# Effect of synaptic cell-to-cell transmission and recombination on the evolution of double mutants in HIV

**DOI:** 10.1101/746131

**Authors:** Jesse Kreger, Natalia L. Komarova, Dominik Wodarz

## Abstract

Recombination in HIV infection can impact virus evolution *in vivo* in complex ways, as has been shown both experimentally and mathematically. The effect of free virus versus synaptic, cell-to-cell transmission on the evolution of double mutants, however, has not been investigated. Here we do so by using a stochastic agent-based model. Consistent with data, we assume spatial constraints for synaptic but not for free-virus transmission. Two important effects of the viral spread mode are observed: (i) For disadvantageous mutants, synaptic transmission protects against detrimental effects of recombination on double mutant persistence. Under free virus transmission, recombination increases double mutant levels for negative epistasis, but reduces them for positive epistasis. This reduction for positive epistasis is much diminished under predominantly synaptic transmission, and recombination can in fact lead to increased mutant levels. (ii) The mode of virus spread also directly influences the evolutionary fate of double mutants. For disadvantageous mutants, double mutant production is the predominant driving force, and hence synaptic transmission leads to highest double mutant levels due to increased transmission efficiency. For advantageous mutants, double mutant spread is the most important force, and hence free virus transmission leads to fastest invasion due to better mixing. For neutral mutants, both production and spread of double mutants are important, and hence an optimal mixture of free virus and synaptic transmission maximizes double mutant fractions. Therefore, both free virus and synaptic transmission can enhance or delay double mutant evolution. Implications for drug resistance in HIV are discussed.

---

**V**irus evolution *in vivo* is a central characteristic of human immunodeficiency virus (HIV-1) infection [20, 14, 31]. Viral evolutionary processes have been shown to drive disease progression through a variety of mechanisms, including evolution of immune escape or evolution towards virus strains with faster replication kinetics, increased cytopathicity, and broader cell tropism [20]. A relatively fast mutation rate of HIV-1 [30], together with a high turnover of the virus during the chronic phase of the infection [41, 15, 37], certainly contributes to the generation and emergence of mutants that drive this disease. These mutational processes are also implicated in the evolution of drug resistance during anti-viral therapy.

In addition to mutations, another mechanism that contributes to virus evolution is recombination [29, 19, 33]. HIV is a diploid virus containing two copies of genomic RNA. If cells are infected simultaneously by different virus strains [29], two different viral genomes can be packaged into the same virus particle. When this virus infects a new target cell, recombination between these two genomes can occur during reverse transcription, when the viral DNA is generated. Recombination has the potential to bring two separate point mutations together in a single virus genome that previously were present in different genomes. Recombination has been shown experimentally to play an important role in HIV-1 infection [33, 32] in situations where the accumulation of two or more mutations is required to achieve a given phenotypic effect. Examples are the generation of virus mutants that are simultaneously resistant against two or more drugs, or mutants that have escaped two or more immune cell clones.

The process of recombination requires the infection of cells with two or more viruses that are genetically different [29]. In HIV, the multiple infection of cells has been shown to be promoted by direct cell-to-cell transmission of the virus, through the formation of virological synapses [16, 7, 1, 39]. Many viruses are transferred simultaneously from the source cell to the target cell, several of which can successfully integrate into the new host cell, making this an efficient mode of infection. Further, experiments have shown that if cells are infected with two distinct virus strains, synaptic transmission promotes the repeated co-transmission of these different strains from one cell to the next [8, 28], which can promote the occurrence of recombination. This was demonstrated both *in vitro* [8] and *in vivo* [28] using HIV-1 infection of humanized mice. *In vivo* data also suggests that the process of synaptic transmission is spatially restricted, meaning that transmission likely occurs to neighboring target cells [28].

The effect of viral recombination on the *in vivo* evolution of HIV has been investigated with mathematical models, revealing a wealth of results, in particular in the context of drug resistant viruses. In [12], recombination was found to be detrimental to the doubly-resistant virus. In [5], the role of recombination was reported to depend on the relative fitness characteristics of single and double mutants, but for most plausible scenarios it was established that recombination slowed down the evolution of resistance. In the models of [2, 6], it was determined that recombination was beneficial for double mutants. In [27] it was clarified that the results strongly depend on the model formulation. In particular, a distinction was made between (i) population genetic (constant population) and population dynamic models, and (ii) stochastic and deterministic models. The model employed in [27] combines a population dynamic description with stochasticity, and finds that recombination decelerates the emergence of drug resistance.

In the present paper we focus on the evolutionary dynamics of double mutant evolution in HIV infection, and how this is influenced by the mode of virus spread (synaptic vs. free virus transmission) and the occurrence of recombination. Just as in [27], we use a stochastic, population dynamic model. In contrast to the above paper, however, we do not use a combined model where “pre-treatment” and “treatment” regimes are both included, but instead focus in more general terms on disadvantageous, advantageous, and neutral mutants. We consider fitness landscapes that range from maximal positive to maximal negative epistasis, expressed by a parameter that ranges from zero to one. Times to double mutant invasion and the fraction of double mutants at defined time points are recorded in the presence and absence of recombination, and for a variety of different virus transmission strategies that range from 100% synaptic to 100% free virus transmission.

## 1 Modeling virus evolution

### 1.1 Stochastic modeling of spatially restricted synaptic virus transmission

Virus dynamics can be modeled by using ordinary differential equations (ODEs) [34, 36]. Extensions of those models that include both free virus and synaptic transmission modes, as well as multiple infection have been since investigated, see [26, 23, 24, 3, 11, 10, 9].

*In vivo* data from humanized mice indicate that synaptic transmission results in spatial clusters of infected cells [28]. In order to explicitly include spatial dynamics of cell-to-cell transmission, we turn to a stochastic agent-based model. This includes both free virus and cell-to-cell transmission, and is adaptable to make either transmission process spatial or non-spatial. We consider a *𝒩 × 𝒩* two-dimensional grid, where each grid point can be empty, contain an uninfected cell, or contain an infected cell. Infected cells can contain any natural number of virus copies. For each time step we randomly make *𝒩*^2^ updates to the grid according to the following rules:

- empty grid points can become uninfected cells with probability *λ*;
- uninfected cells can die with probability *d*;
- infected cells can die with probability *a*, infect another cell by free virus transmission with probability *β*, or infect another cell by cell-to-cell transmission (with *S* copies of the virus) with probability *γ*. During the infection processes, a target spot is chosen randomly either from the entire grid or the local neighborhood. If that target spot contains a susceptible cell (uninfected or already infected), the infection event proceeds, otherwise it is aborted.

We assume that synaptic transmission can only occur to one of the eight nearest neighbors, while free virus transmission can occur to any cell on the grid. Basic simulations of this model can be seen in Supplemental figures S3 and S4.

It is straightforward to extend the agent-based model to include two virus strains that compete for the same target cell population (also see corresponding ODEs (8-9) of the Supplement, Section 1). In this setting, a cell can be infected by *i* copies of virus strain A and *j* copies of virus strain B. If a cell containing both virus strains is chosen for an infection event, the probability to transmit a given virus strain is proportional to the fraction of the strain among all viruses in the cell if the two strains are neutral with respect to each other. If the two strains have different replication rates, the fitness difference is implemented during the infection event, which can correspond to different rates of reverse transcription. That is, an infecting strain is again chosen randomly with a probability that is proportional to its fraction in the cell. A disadvantageous / advantageous mutant would then have a lower / higher probability to infect the chosen target cell upon infection. In the neutral case, drift is observed with the eventual fixation of one of the virus strains. If the two virus strains have different fitness, the strain with the larger basic reproductive ratio [34] wins. Both of these cases can be seen in Supplemental figure S3.

### 1.2 Mutations and recombinations

We consider a virus population that can mutate at two different sites, denoted by *a* and *b*. Simulations are started with unmutated wild-type cells, *ab*. Single-mutant viruses (*Ab* or *aB*) can be generated during infection by point mutations, which occur with a probability *µ* per site. Each single-mutant can in turn mutate further to give rise to a double mutant *AB*. Note that a wild-type virus can directly mutate into a double mutant with a probability *µ*^2^ if both sites mutate during the same reverse transcription event. The model also takes into account back-mutations, which again occur with a probability *µ* during an infection event. All the possible mutation events are illustrated in figure S1 of the Supplement.

Apart from mutations, however, a double mutant can also be generated through the recombination of different single-mutant viruses. This is implemented as follows. When viruses from a given source cell are chosen to infect a target cell, two virus genomes are randomly chosen with a probability that is proportional to the fraction of their abundance in the cell. The first virus genome that is chosen is the template from which reverse transcription is initiated. If no recombination occurs, reverse transcription is assumed to proceed on this genome only. Recombination is assumed to occur with a probability *ρ*. In this case, the reverse-transcribed virus is assumed to be a recombinant, the identity of which depends on the two infecting genomes. Figure S2 of the Supplement list all recombination events that can occur.

There are two recombination processes in particular that are important: (i) *Ab* + *aB → AB* with probability *ρ/*2 (or *ab* with probability *ρ/*2), and (ii) *ab* + *AB → Ab* with probability *ρ/*2 or *aB* with probability *ρ/*2. These processes capture two roles of recombination that have been previously discussed in the literature [5]. Recombination between two single mutants can promote the generation of the double mutant, but recombination can also break up a double mutant upon recombination with the wild-type virus.

### 1.3 Simulations of the model and parameter values

We initialize the infection by randomly and uniformly spreading an equilibrium number of infected cells across the grid. These cells are singly infected with the wild type. We used a mutation rate of 3 × 10^*-*5^ [30] and a recombination rate *ρ* = 0, 0.1, 0.2, and 0.5. Most of the parameters of this system are unknown. The average life-span of productively infected cells is around 2 days^1^ [37], and the basic reproductive ratio of HIV (and SIV) has been estimated to be around 8 [38, 35]. The unknown parameters were chosen arbitrarily such that the basic reproductive ratio of the virus was between 5 and 8 (depending on assumptions on mutant fitness), and are provided in the figure legends. In regimes where the basic reproductive ratio of the virus is around 8, the average multiplicity of infection in cells lies between 4-14, depending on how prevalent synaptic transmission is assumed to be. Widely varying estimates for average infection multiplicities have been published [19, 17, 18, 40], and there is some uncertainty about that. To investigate scenarios in which the average infection multiplicity is on the lower end (between 1-3, depending on viral transmission mode), we modified the model to track time since infection and assumed that the probability of superinfection declines over time due to receptor down-modulation [11]. This is described in the Supplementary Materials (Section 4).

To investigate the relative contribution of free virus transmission (*β*) and cell-to-cell transmission (*γ*) we ran the model for different combinations of *β* + *γ* = *c*, where *c* is a fixed constant, ranging from purely synaptic to purely free virus transmission. The average outcome of the simulations were determined, including the average generation rate of double mutants, the average fraction of double mutants at a specific time point, and the time until the double mutant population grew to 90%.

## 2 Generation and spread dynamics of the double mutant

We will present all results for a range of transmission mode combinations, ranging from 100% synaptic transmission to 100% free virus transmission. In this section, however, we will mainly discuss under what fitness landscapes and assumptions recombination generally promotes the presence of double mutant populations, and when it works against them. The subsequent section will then discuss in more detail how these basic patters are modulated by synaptic versus free virus transmission.

### 2.1 Fitness landscapes and epistasis

Our investigation will span a variety of fitness landscapes, including neutral, advantageous, and disadvantageous mutants. Let us assume that a mutation in site A or B results in an identical change in the fitness of the virus. Then, possible fitness landscapes can be separated into three groups for both advantageous and disadvantageous mutants [5]: negative epistasis, no epistasis, and positive epistasis, see figure 1(a) for examples of these.

**Figure 1:**
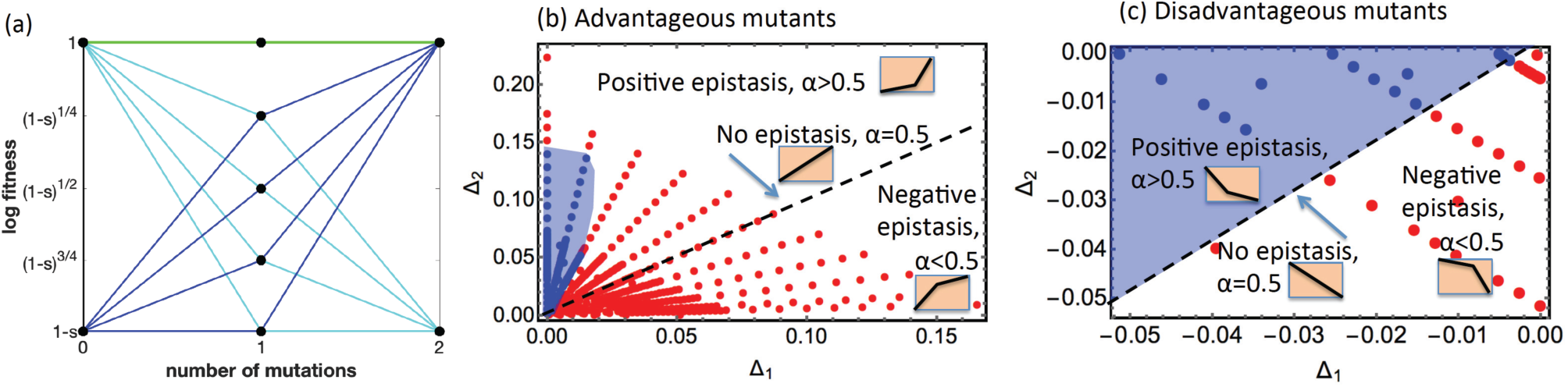
Summary of different scenarios. (a) Examples of fitness landscapes used in the simulations neutral (green), disadvantageous (cyan), advantageous (dark blue). Without epistasis the single-mutant fitness is given by *f* = (1 − *s*)^1/2^. For negative and positive epistasis examples in the figure, it is given by (1 − *s*)^1/4^ and (1 − *s*)^3/4^ respectively. For the extreme form of positive epistasis, single-mutant fitness is the same as that of wild types, 1*-s*. (b-c) Role of recombination for different fitness landscapes. The horizontal axis is Δ_1_ and the vertical axis is Δ_2_, which are the relative log fitness values of single and double mutants, respectively. Each of the dots corresponds to a particular fitness landscape. (b) Advantageous mutants: Red dots correspond to runs in which recombination accelerated double hit mutant invasion to 90%, while blue dots indicate that recombination slowed down invasion. (c) Disadvantageous mutants: Red dots indicate that recombination increased the double mutant fraction at *T* = 10^5^, while blue dots mean that recombination reduced the double mutant fraction. Blue shading marks the regions where recombination suppresses double hit mutants. The dashed black line corresponds to the cases of no epistasis (*α* = 0.5) and separates the regions with positive epistasis (*α >* 0.5) and negative epistasis (*α <* 0.5). For both (b) and (c), we fixed the probability of free-virus transmission at 40% (*β* = 0.04); the rest of the parameters are as in figure 2. The determination on whether recombination suppressed or enhanced double hit mutants was made by a statistical comparison of the averages over many runs, using the t test.

Notice that each of the landscapes with advantageous mutants can be written as a triple of numbers,

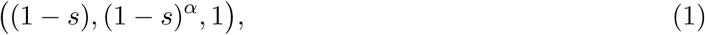

which represent fitness values of the wild types, one hit mutants and double mutants respectively. Here *s >* 0 measures the amount of advantage, and *α* represents epistasis. We have *α >* 1/2 for positive epistasis, *α <* 1/2 for negative epistasis, and *α* = 1/2 for no epistasis landscapes. Define the relative (log) fitness of the one-hit mutants compared to that of wild types, Δ_1_ = ln(1 − *s*)^*α*^ − ln(1 − *s*) = (1 − *α*)*|* ln(1 − *s*)*|*, and the relative (log) fitness of the two-hit mutants compared to that of one-hit mutants, Δ_2_ = ln(1) − ln(1 − *s*)^*α*^ = *α|* ln(1 − *s*)*|*. Note that the sum of the two coordinates, Δ_1_ + Δ_2_ = *|* ln(1 − *s*)*|* represents the relative log fitness of the two-hit mutants compared to the wild types.

Similarly, each of the landscapes with disadvantageous mutants presented in figure 1(a) can be written as a triple of numbers,

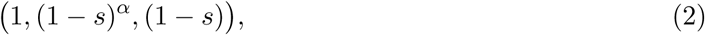

where *s >* 0 measures the amount of disadvantage. The relative (log) fitness of the one-hit mutants compared to that of wild types, Δ_1_ = ln(1*-s*)^*α*^ − ln(1) = *α* ln(1*-s*). The relative (log) fitness of the two-hit mutants compared to that of one-hit mutants, Δ_2_ = ln(1 − *s*) − ln(1 − *s*)^*α*^ = (1*-α*) ln(1 − *s*). Again, the sum of the two coordinates, Δ_1_ + Δ_2_ = ln(1 − *s*), represents the relative log fitness of the two-hit mutants compared to the wild types.

### 2.2 Advantageous mutants

A reasonable measure of double mutant success is the time it takes for the double hit mutant to reach 90% of all infected cells. The following factors trade-off to determine whether recombination boosts or suppresses double mutant spread: (i) Recombination between single mutants increases the rate at which double mutants are generated; (ii) recombination between double mutants and wild-type can break apart double mutants. (iii) the strength of selection of the double mutant defines how long the previous two factors are at play.

The net effect of recombination depends on the degree of the selective advantage, parameter *s*. For stronger advantages (larger values of *s*), recombination reduces the time to double mutant invasion (figure 2(b)). For lower selective advantages (lower *s*), however, recombination increases the time to double mutant invasion (figure 2(a)). The stronger the selective advantage, the quicker the double mutants spread at the expense of the wild-type, and then less likely it is that detrimental recombination events with the wild-type virus occur. This is illustrated with specific realizations of the stochastic dynamics in the Supplementary Section 3, figure S6.

**Figure 2:**
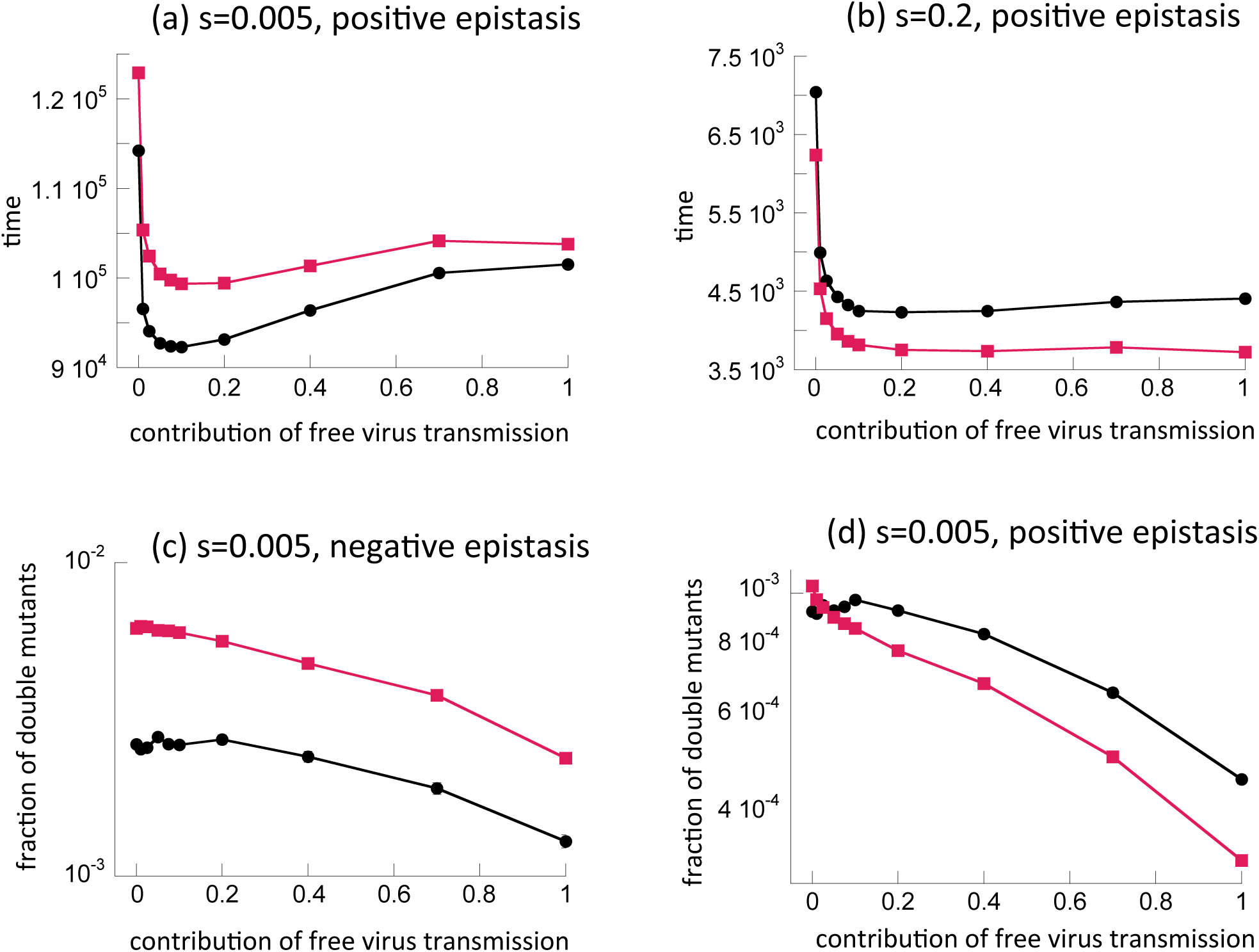
The role of recombination in (a,b) advantageous and (c,d) disadvantageous mutant dynamics. Red: with recombinations, and black: without recombination. (a-b) The time until the advantageous mutant reaches 90%, as a function of the fraction of free virus transmission. The means and standard errors are shown. (a) *s* = 0.005, *α* = 0.75. (b) *s* = 0.2, *α* = 0.75. (c-d): The fraction of disadvantageous mutants at time *T* = 10^5^, as a function of the fraction of free virus transmission. The means and standard errors are shown. (c) *s* = 0.005, *α* = 0.25 (d) *s* = 0.005, *α* = 0.75. The parameters are: *β* + *γ* = 0.1, *S* = 3, *λ* = 1, *d* = 0.01, *a* = 0.02, *𝒩* = 100, *µ* = 3 × 10^*-*5^. All averages are based on at least 10^4^ simulations.

The selective advantage threshold below which recombination slows double mutant invasion depends on the nature of the fitness landscape, in particular the value of *α*. This is summarized in figure 1(b). The horizontal axis is Δ_1_ (fitness difference between single-mutant and wild-type) and the vertical axis is Δ_2_ (fitness difference between double and single mutants). Each point in this coordinate system corresponds to a unique fitness landscape. The red color means that recombination events promote double mutant invasion, and blue means that they suppress this process. This picture has been composed by assuming 40% free virus transmission, and 60% synaptic transmission. Arrays of points radially fanning out of the origin correspond to landscapes with the same level of epistasis (the same value of *α*) but different selection strength (the closer to the origin, the lower *s*). We observe that for any level of epistasis, for sufficiently high fitness advantage, recombinations are advantageous for double hit mutants. As we decrease fitness *s*, however, there comes a point where recombinations no longer enhance double mutants but instead suppress them. In other words, any radial line will enter the blue region if it is sufficiently close to the origin. Recombinations can suppress the double mutant population significantly even for relatively large fitness advantages (large value of *s*) if *α* is relatively large and converges to one, i.e. for large positive epistasis (points close to the vertical axis in figure 1(b)). For lower values of *α* (weaker positive epistasis, no epistasis and negative epistasis), however, the transition happens for progressively smaller values of *s*. For large negative epistasis, the transition happens for very small values of *s* (for example, calculations show that for *α* = 0.25, the blue region starts at about *s ≈* 10^*-*5^, which is too small to see clearly in the figure and irrelevant for practical purposes, because such mutants are effectively neutral and take on average very long times to rise). The intuitive explanation for these observations is that lower values of *α* result in a more pronounced fitness advantage of single-hit mutants compared to wild-type virus. This in turn results in a faster exclusion of the wild-type virus population, and thus reduces the chances that recombinations break the double mutants. Hence, the parameter regime in which recombinations have a net negative effect on the double mutant population becomes more restrictive.

### 2.3 Disadvantageous mutants

A selective disadvantage leads to competitive exclusion in the absence of mutational processes. In the presence of mutational processes, disadvantageous mutants on average persist at an equilibrium level determined by the balance between mutation and selection. Hence, we determined the average fraction of double mutants at a time when this equilibrium has been reached, for different combinations of synaptic and free virus transmission (see Supplement Section 3.2 for details).

Recombination increases the double mutant population at the selection-mutation balance for negative epistasis (figure 2(c)), but tends to reduce it for positive epistasis if a sufficient amount of free virus transmission is assumed to occur (figure 2(d)). Similar results have been reported in the context of HIV drug resistance evolution [27]. If most virus transmission, however, occurs through the synaptic route, figure 2(d) suggests that the opposite becomes true: Now, recombination can increase the mutant levels for positive epistasis as well. This will be explored in more detail below.

These trends are further illustrated in figure 1(c), assuming that a mixture of free virus and synaptic transmission occurs: As the parameter *α* is increased, the effect of recombination on the equilibirium level of double mutants changes from beneficial to detrimental. For the particular mixture of synaptic and free virus transmission chosen in this figure, recombination increases double mutant levels for negative epistasis (red region), and suppresses double mutant levels for positive epistasis (blue region). An increase in the parameter *α* results in a lower fitness of single mutants relative to the wild-type virus. This results in a higher prevalence of the wild-type virus, and thus in higher chances for the wild-type to recombine with and break apart the double mutant; see Supplement Section 3.3 for an ODE approximation of these results.

### 2.4 Neutral mutants

It follows from the above analysis that recombination delays the drift of neutral mutants towards dominance. Consider very weakly advantageous mutants in figure 1(b). We can see that the origin is contained in the blue region, that is, as the selective advantage *s* → 0, we expect recombinations to delay the rise of double mutants.

For neutral mutants, however, the rise to dominance will take a very long time. Interestingly, different results are obtained if we look at the fraction of double mutants at a relatively early time point *T*. Figure 3(a) shows that recombination increases the fraction of double mutants at time *T*. This can be understood by considering the early vs long-term dynamics of neutral mutants. In the long-run, the populations will converge to a state where all four virus strains fluctuate around comparable fractions. This steady state is the same whether recombination occurs or not. The speed with which the double mutant rises towards this steady state, however, is influenced by the occurrence of recombination (figure 3(b,c)). Initially, the populations of single mutants are generated by mutations and rise by drift. In the absence of recombinations, double mutants are created and destroyed by mutations and also experience drift (panel (b)). In the presence of recombinations, however, double hit mutants initially enjoy positive selection due to relatively frequent recombination events between complimentary single hit mutants (which greatly outweigh the “breaking” recombination events of the double mutants with the wild type, due to the low levels of the former population). This can be seen in panel (c) of figure 3. Once the levels of double hit mutants increase, however, the “making” and “breaking” recombination events begin to balance each other and the dynamics return to neutral.

**Figure 3:**
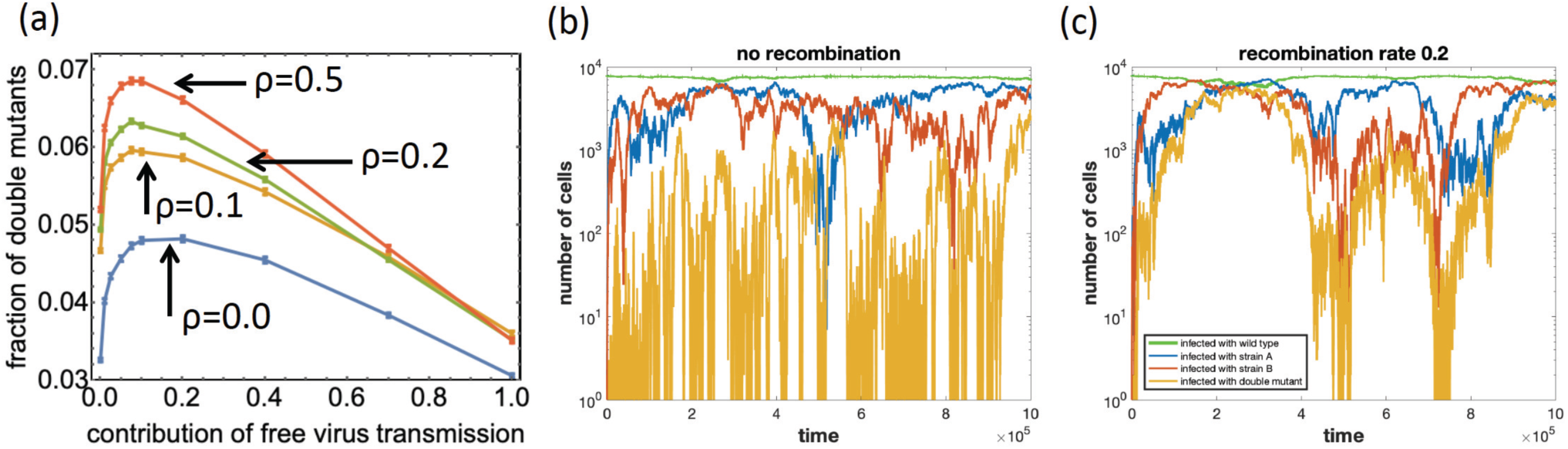
Neutral mutants. (a) The fraction of mutants after *T* = 10^5^ steps. The horizontal axis is the fraction of free virus transmission. 4 values of *ρ* are presented from *ρ* = 0 (no recombinations) to *ρ* = 0.5 (maximal recombinations). The means of at least 40, 000 runs at each location and standard errors are shown. (b,c) The dynamics of cell populations, typical time-series: (b) no recombinations, (c) with recombinations; we used *β* + *γ* = 0.1. The other parameters are: *S* = 3, *λ* = 1, *d* = 0.01, *a* = 0.02, *𝒩* = 100, *µ* = 3 × 10^*-*5^, *ρ* = 0.2.

## 3 Mode of viral transmission and the effect of recombination on double mutant populations

The last section examined under what fitness landscapes recombination promotes or hinders the existence of double mutants. For advantageous and neutral mutants, these results remain robustly independent of the mode of virus transmission (Supplementary Section 5.1). For disadvantageous mutants, however, we noted that results can change if most virus transmission is assumed to be synaptic. Figure 2 showed that while for smaller values of *α* (negative epistasis, panel (c)), recombination lead to an increase in double mutant levels, for large values of *α* (positive epistasis, panel (d)), the opposite occurred and recombination reduced the double mutant levels. At the same time, however, figure 2(d) indicated that if most virus transmission occurs through the synaptic pathway, recombination remains helpful for the double mutant population even for positive epistasis. This is explored in more detail in figure 4, which plots the equilibrium level of a disadvantageous mutant as a function of the parameter *α* for both extreme transmission modes: 100% free virus and 100% synaptic. If only free virus transmission occurs (panel (a)), recombination increases the double mutant fraction for *α <* 0.5 (negative epistasis), while it decreases it for *α >* 0.5 (positive epistasis). In contrast, if only synaptic transmission occurs (panel (b)), recombination always increases the number of double mutants, regardless of the value of *α*, although the double mutant levels in the presence and absence of recombination become practically indistinguishable for large values of *α* (strong positive epistasis). Supplemental figure S11 contains further simulations showing the robustness of these patterns for different levels of mutant disadvantage. Similar patterns hold for lower infection multiplicities (Section 4, Supplementary Materials). While for lower multiplicities, the equilibrium fraction of double mutants can still be slightly reduced by recombination for purely synaptic transmission, this reduction is much less than in the presence of only free virus transmission, thus confirming the protective effect of synaptic transmission even in the low multiplicity scenario. Therefore, if positive epistasis is present, as is suggested for drug resistance mutations in HIV [4], a prevalence of synaptic transmission can protect against the negative effects of recombination on the level at which drug-resistant mutations pre-exist before the start of treatment.

**Figure 4:**
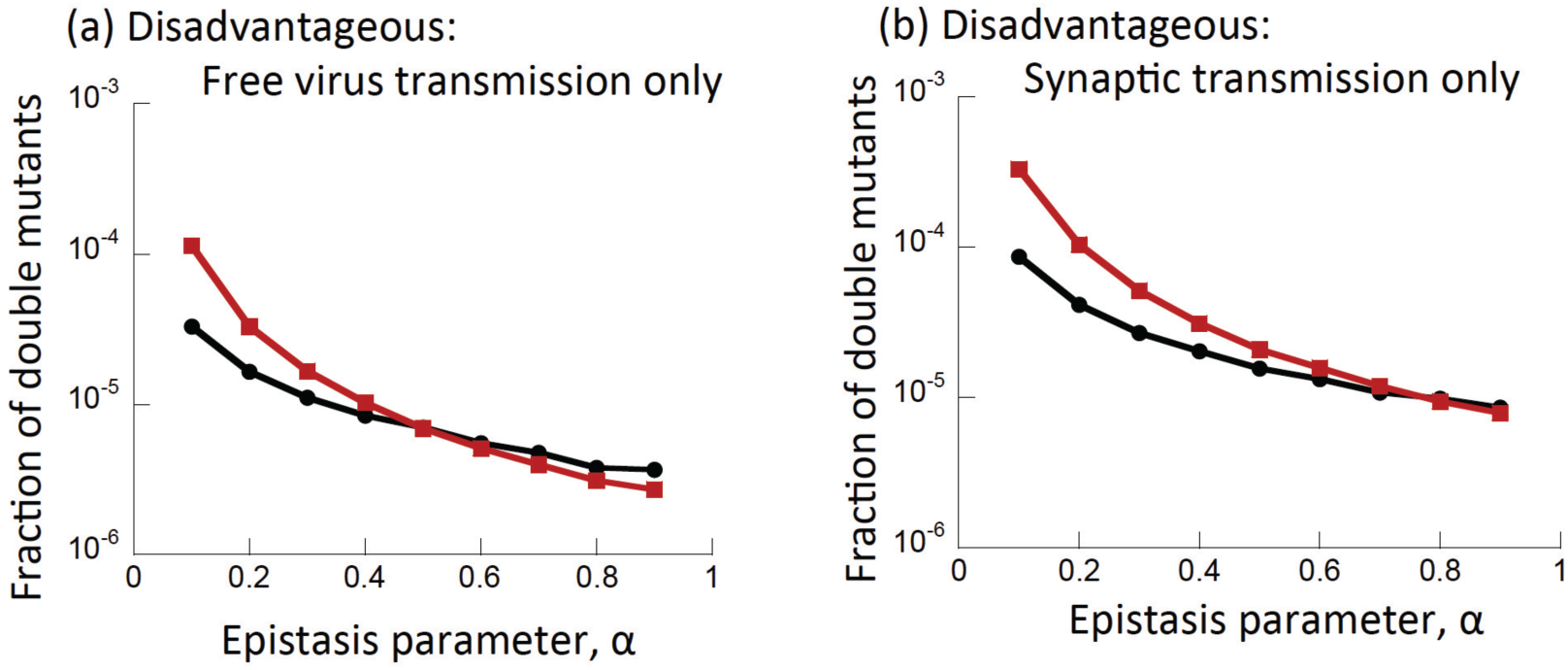
The role of recombinations under different transmission modes, for disadvantageous mutants. Shown is the temporal average of the fraction of double mutants at selection-mutation balance, as a function of that parameter *α*, defining the nature and extent of epistasis. Red denotes simulations with recombination and black without recombinations. (a) Free virus transmission only, (b) synaptic transmission only. *s* = 0.05, and other parameters are as in figure 2. Standard errors are too small to see.

The intuitive explanation for the detrimental effect of recombination on the double mutant population at larger values of *α* was given in the previous section: For larger values of *α*, the fitness of single mutants relative to wild-type viruses becomes lower. This leads both to a slower rate of double mutant production, and to a higher prevalence of wild-type viruses that can recombine with the double mutant and break it. If most of virus transmission occurs through virological synapses, however, the spatially restricted virus spread that is assumed to occur with this transmission mode results in the generation of single and double mutant “clusters” or “islands”. Single-mutant islands protect them from being outcompeted, resulting in larger numbers and thus a higher rate of double mutant generation. Double mutant islands isolate them from contact with wild-type virus, which prevents those detrimental recombination events from occurring. These dynamics are similar to the effect of “mutant islands” discussed in [21, 25].

## 4 Mode of viral transmission and the rate of double mutant emergence

All simulations were performed for varying combinations of synaptic and free virus transmission, yet we have so far not discussed the effect of this itself on the emergence of the double mutant population. A number of factors trade off to determine what combination of synaptic and free virus transmission is optimal for the double mutant population. On the one hand, synaptic transmission results in the simultaneous transfer of multiple viruses from the source cell to the target cell, which increases the rate at which mutants are generated, and increases the rate of co-transmission of genetically different viruses, which in turn promotes the occurrence of recombination. On the other hand, if synaptic transmission is spatially restricted, as indicated by data [28], the rate at which the number of infected cells increases is slower under this mode of transmission, and it is less likely that genetically different strains come together in the same cell. For the different mutant types, the net effect is as follows:

### Disadvantageous mutants

For disadvantageous mutants, more synaptic transmission tends to increase the equilibrium levels of double mutants at the selection-mutation balance (figure 2(c,d) and Supplemental figure S9(a,b) for the model with limited multiplicity). The main driving force responsible for the abundance of double mutants is production. This is maximized by synaptic transmission, because under this mechanism, there are more possibilities for mutations. Spread to higher levels is not an important force for disadvantageous mutants.

### Advantageous mutants

In the case of advantageous mutants, the rate of double mutant invasion tends to be increased by free virus transmission, and purely synaptic transmission results in the slowest rate of invasion (figure 2(a,b)). The reason is that in this scenario, the spread of the double mutant from low to high numbers is the driving process, and this is slower for synaptic transmission, which is assumed to be spatially restricted [28]. While increasing the contribution of free virus transmission generally speeds up mutant invasion, this trend can weaken or reverse for larger fractions of free virus transmission, which can result in a shallow optimum, see figure 2(a,b) and Supplemental figure S9(c-e) for the model with limited multiplicity. The reason is that in the absence of significant synaptic transmission, fewer overall infection, and hence reverse transcription, events occur, which delays mutant production.

### Neutral mutants

In the neutral case, and also for very weakly advantageous and disadvantageous mutants, a mixture of both free virus and cell-to-cell transmission maximizes the fraction of cells infected with the double mutant (figure 3(a) and Supplemental figure S9(f) for the model with limited multiplicity). This result is similar to what was observed in Section 2 of the Supplement, where the generation time of double hit mutants was studied. Here we observe that this holds even in the absence of recombination, and is more pronounced. The reason is that while more synaptic transmission results in the simultaneous transfer of multiple viruses, and hence in more chances to mutate, it also slows down the increase of the infected cell population due to the assumed spatial restriction. In this scenario, both production and spread play important roles.

## 5 Discussion and Conclusion

We aimed to comprehensively analyze the effect of recombination on double mutant evolution in the context of HIV, depending on the details of fitness landscapes and the assumptions about the mode of viral spread (relative importance of synaptic versus free virus transmission). This is different from previous approaches, which focused more specifically on the evolution of drug resistance in HIV in the context of only free virus transmission, and concentrated on specific fitness landscapes characterized by positive or negative epistasis. Our approach characterized the fitness landscape by the parameter *α*, which could be continuously varied from 0 to 1, thus capturing all fitness landscapes ranging from negative to positive epistasis for advantageous and disadvantageous mutants. The constraint in our fitness landscapes was that the two different single-hit mutants were assumed to have identical fitness.

The opposing effect of recombination to make and break double mutants played out as follows in the model analyzed here: For advantageous mutants, recombination largely accelerates double mutant invasion except for cases of very strong positive epistasis with an intermediate fitness advantage of the mutants, or in cases where the fitness advantage becomes relatively low. The mode of viral spread does not modulate these patterns. If the mutants are disadvantageous, however, the mode of virus spread can significantly influence the effect of recombination on the equilibrium level of double mutants at selection-mutation balance. If the contribution of free virus transmission to virus spread lies above a threshold, recombination increases the double mutant population for negative epistasis, but decreases it for positive epistasis. If the dominant mode of virus spread is synaptic transmission, however, the negative effect of recombination for positive epistasis is greatly reduced, indicating a protective effect on the persistence of disadvantageous double mutants. In fact, for higher multiplicities, recombination increases double mutant levels even for positive epistasis if synaptic transmission is the dominant mode of virus spread. Finally, for neutral mutants, we observed that recombination always delays the rise of double mutants to dominance, but at the same time increases double mutant fractions measured at a relatively early time points in the dynamics. Interestingly, neutral double mutant dynamics in the presence of recombination are characterized by an “advantageous” initial growth phase before converging to neutral drift, which explains the positive effect of recombination on early double mutant fractions.

These findings have implications for the pre-existence of multi-drug resistant HIV mutants before the start of therapy. In the absence of treatment, resistant mutants typically carry a fitness cost. Moreover, evidence for positive epistasis has been observed in HIV resistance evolution [4]. Therefore, a relatively high rate of synaptic transmission could significantly increase the chances that multi-drug resistant virus mutants are present at selection-mutation balance before treatment is initiated.

The way in which the mode of virus spread was observed to influence the rate of double mutant emergence was driven by two opposing effects: synaptic virus transmission increases double mutant production in our model, but slows down double mutant spread due to the experimentally supported assumption that synaptic transmission is associated with spatially clustered dynamics [28]. For disadvantageous mutants, production is the main driving force, and hence purely synaptic transmission results in highest mutant levels. For advantageous mutants, double mutant spread is a crucial factor, and hence, free virus transmission tends to speed up mutant invasion in our model. For neutral mutants, both production and spread are similarly important, and hence, there is an optimal combination of free virus and synaptic transmission that maximizes double mutant fractions.

There is some controversy in the literature about the average multiplicity of infected cells in vivo. While some papers reported significant levels of multiple infection, especially in tissue compartments [19, 40], other publications found an infection multiplicity close to one, both in the blood and tissues [17, 18]. Reasons for the discrepancy could be the methodology that was used to measure multiplicity, and also the T cell subsets that were taken into account during this analysis. In the light of data that document an important contribution of recombination to the *in vivo* evolution of HIV [29, 19, 33, 32], it is likely that a sufficient amount of multiple infection occurs. An important role of multiple infection is further suggested by studies that document a very efficient infection process during synaptic cell-to-cell transmission, resulting in the simultaneous transfer of multiple viruses from the source cell to the target cell [7, 16, 39]. Further, the frequent co-transmission of different virus strains was observed both *in vitro* and *in vivo* [8, 28]. An important point in our analysis was that a model with reduced infection multiplicity due a declining ability to super-infect over time resulted in similar insights.

Another crucial assumption of our model was that synaptic cell-to-cell transmission was characterized by spatially restricted virus spread. While imaging studies have shown an ability of immune cells to move about within tissues [13], our work on humanized mice demonstrated that virus spread in the presence of synaptic, cell-to-cell transmission, was characterized by the spatial clustering of infected cells [28], which supports the assumption we made. If it were assumed that synaptic transmission follows mass action law, then several results would change, since synaptic transmission would no longer give rise to slower virus spread than free virus transmission. A certain amount of cell migration could be incorporated into the model, and this would be an interesting future extension of the current work.

The relative contribution of synaptic and free virus transmission to virus spread *in vivo* is still not well-understood. *In vitro* experiments have estimated that the two transmission modes contribute approximately equally to virus spread [22], but conditions *in vivo* are likely significantly different, and this could have a large impact on these dynamics. Our results indicate that both free virus and synaptic transmission have important and different effects on double mutant populations, depending on the nature of the mutants. Free virus transmission promotes the invasion of advantageous double mutants, while synaptic transmission promotes the existence of disadvantageous double mutants at selection-mutation balance. Further, we observed synaptic transmission to protect against negative effects of recombination for disadvantageous double mutants characterized by positive epistasis. These selective forces likely shape the balance between synaptic and free virus transmission towards which HIV has evolved.

## Acknowledgements

This work was supported by NSF grant DMS 1662146 /1662096.

## Footnotes

1 Note that the death rate parameter used in our simulations corresponds to arbitrary time units, and the 2 days life span is recovered by an appropriate scaling of the time units.

