## Supplementary Information for "Effect of synaptic cell-to-cell transmission and recombination on the evolution of double mutants in HIV"

### 1 Deterministic and stochastic modeling of synaptic and free virus transmission

#### 1.1 Deterministic modeling

Here, we review a previously published mathematical modeling framework to study the role of synaptic and free virus transmission in HIV dynamics [4, 2, 3]. This modeling approach is based on ordinary differential equations, which means that perfect mixing of populations occurs with both transmission modes, i.e. synaptic transmission is not spatially restricted. Let $x_i(t)$ be the population of cells infected with $i$ copies of the virus at time $t$. Let $\gamma_j^m$ be the probability for a cell infected with $m$ viruses to transmit $j$ viruses per synapse. Let $N$ be the maximum number of copies of the virus that a single cell can contain, known as the infection multiplicity. The model equations are

$$\dot{x}_0 = \lambda - dx_0 - \sum_{m=1}^{N} x_m \sum_{j=1}^{N} \gamma_j^m x_0, \tag{1}$$

$$\dot{x}_i = \sum_{m=1}^{N} x_m \Big( \sum_{j=1}^{i} \gamma_j^m x_{i-j} - x_i \sum_{j=1}^{N-i} \gamma_j^m \Big) - ax_i, \tag{2}$$

where $\lambda$ is the constant production rate, $d$ is the death rate of uninfected cells, and $a$ is the death rate of infected cells. Cell-to-cell (synaptic) transmission is given by the $\gamma_j^m$ parameters. Free virus transmission (set $j = 1$) is implicitly included. Since both processes are non-spatial, in the model given by equations (1-2), free virus transmission and cell-to-cell transmission are very similar. The only difference is that during a cell-to-cell transmission event, $j$ copies of the virus can be passed, whereas during a free virus transmission event only one copy of the virus can be passed. This model assumes that kinetic parameters are independent of infection multiplicity, because there is currently no evidence to the contrary.

This model is characterized by two outcomes / equilibria. The disease-free equilibrium is given by

$$x_0 = \frac{\lambda}{d}, \tag{3}$$

$$x_i = 0 \text{ for } 1 \le i \le N. \tag{4}$$

Virus persistence is described by the following equilibrium expressions:

$$x_0 = \frac{a}{\beta}, \tag{5}$$

$$z = \frac{\lambda}{a} - \frac{d}{\beta}, \tag{6}$$

where $z$ denotes the sum of all infected cell sub-types. Additionally, we can calculate the steady state value for each individual infected population. We have from [4] that if the probability to transmit the virus does not depend on how many viruses the cell contains, then each individual infected population is given by

$$x_i = \frac{\sum_{s=1}^{i} \gamma_s x_{i-s}}{\tilde{a} + p_i} \text{ for } 1 \le i < N. \tag{7}$$

Here, $\tilde{a} = \frac{a}{z}$ and the probability to accept virus particles, $p_i$, is given by $p_i = \sum_{x=1}^{N-1} \gamma_s$. As done in [4], an explicit solution for $x_i$ can also be constructed.

An important measure in virus dynamics is the basic reproductive ratio of the virus, $R_0$, denoting the average number of newly infected cells produced by a single infected cell when placed into a pool of susceptible cells [6, 7]. In a deterministic model, if $R_0 > 1$, the virus successfully establishes an infection, and if $R_0 < 1$, virus extinction occurs. For the model written down here, the basic reproductive ratio of the virus is given by $R_0 = \frac{\beta\lambda}{ad}$. This is the same expression as the one derived from the basic model of virus dynamics in the absence of multiple infection and synaptic transmission [6, 7]. The reason for this is that kinetic parameters, such as the rate of virus production or the rate of virus-induced cell death, are assumed to be independent of infection multiplicity.

As done in [4], this model can be extended to include competition between two or more virus strains. Let $x_{ij}(t)$ be the population of cells infected with $i$ copies of virus strain A and $j$ copies of virus strain B at time $t$. Let $\gamma_{qp}^{mk}$ be the probability for a cell infected with $m$ copies of virus strain A and $k$ copies of virus strain $B$ to transmit $q$ copies of virus strain $A$ and $p$ copies of virus strain $B$. The model equations for competition between two strains are

$$\dot{x}_{00} = \lambda - dx_{00} - x_{00} \sum_{m=0}^{N} \sum_{\substack{k=0 \\ m+k>0}}^{N-m} x_{mk} \sum_{q=0}^{N} \sum_{\substack{p=0 \\ p+q>0}}^{N-q} \gamma_{qp}^{mk}, \tag{8}$$

$$\dot{x}_{ij} = \sum_{m=0}^{N} \sum_{\substack{k=0 \\ m+k>0}}^{N-m} x_{mk} \Big( \sum_{q=0}^{i} \sum_{\substack{p=0 \\ p+q>0}}^{j} x_{i-q,j-p} \gamma_{qp}^{mk} - x_{ij} \sum_{q=0}^{N-i} \sum_{\substack{p=0 \\ p+q>0}}^{N-i-j-q} \gamma_{pq}^{mk} \Big) - ax_{ij}. \tag{9}$$

In equation (9), we assume $i+j > 0$ and $i+j \leq N$. We also assume for all of the double summations that the two indices are not zero simultaneously. These equations can be extended naturally to include competition between any number of additional strains. As shown in [4], the outcome of this competition depends on the relative value of $R_0$ of the two virus strains. If the values of $R_0$ are identical, the strains are neutral with respect to each other, and an infinite number of equilibria exists, depending on the initial conditions. If the two strains have different values of $R_0$, then the strain with the higher $R_0$ wins and excludes the other strain.

##### 1.2 Modeling mutations and recombinations

Figure 1 presents all the (forward and backward) mutation processes that can occur between the four types. Figure 2 presents all possible recombination events that can happen in the presence of two types of mutations. Only two events result in types different from either of the recombining types: the creation of a double hit mutant when mutant $A$ recombines with mutant $B$, and the destruction of a double hit mutant (making it into a single mutant) when it recombines with the wild type virus.

##### 1.3 Spatial stochastic simulations

Figure 3 shows a comparison of the stochastic, agent based model with the deterministic ODE model. The left panel shows numerical solutions of the ordinary differential equations together with the agent based simulation. The numerical solution is represented with dotted lines and a single typical agent based simulation is represented with the solid lines. The different colors represent the number of cells infected with the respective number of copies of the virus. The right panel shows a comparison of agent based simulations for two neutral virus strains (upper right panel) versus two non-neutral strains (lower right panel). In the neutral case, drift is observed with the eventual fixation of one of the virus strains. In the non-neutral case, we assume that strain A has higher fitness over strain B. As a result, strain A will fixate.

Figure 4 shows snapshots of a typical stochastic spatial simulation with infection by a single virus strain. The left panels show a simulation which includes only free virus transmission, whereas the right panels show a simulation which includes only cell-to-cell transmission. The top panels show the status of each grid point, where red grid points denote infected cells. The bottom panels show the multiplicity of infection of each cell, with darker colors representing higher multiplicity. While non-spatial free virus transmission leads to mostly singly infected cells uniformly spread out across the grid, we see here that cell-to-cell transmission leads to spatial clumps of superinfected cells. This is because cell-to-cell transmission leads to the repeated infection of nearby cells, where each time an infection events occurs, multiple copies of the virus are passed.

#### 2 Generation of double mutants

In this section we show that a combination of free virus and cell-to-cell transmission results in faster generation of double mutants by recombination. To study double-hit mutant generation, we ran the simulation repeatedly and recorded the time at which the double mutant was first generated, at which point the simulation was terminated. The average time of double mutant generation for various combinations of synaptic and free virus transmission is shown in figure 5.

In the absence of recombination, double mutant generation occurs fastest with purely synaptic transmission and takes longer as the contribution of free virus transmission is increased. This is because each time a synaptic infection event occurs, multiple viruses are transferred from the source cell to the target cell, thus increasing the number of mutation events that can occur during reverse transcription.

In the presence of recombination, however, we observe that double mutant generation occurs fastest for a mixture of free virus and synaptic transmission (figure 5). The reason is the existence of a tradeoff. Free virus transmission is efficient at bringing together two distinct virus strains (single mutant A and single mutant B) in the same cell, which is essential for recombination (creating the double hit mutant) to occur. Once in the same cell, however, the two virus strains are likely to disperse to different target cells rather than being repeatedly co-transmitted, which limits opportunities for recombination. In contrast, synaptic transmission promotes the co-transmission of two different virus strains once they have come together into the same cell [5], which generates more opportunities for recombination to occur. At the same time, however, the spatial nature of this process makes it less likely that they come together in the same cell to start with. The observed optimum thus presents the best solution to this tradeoff.

The same general trends hold if both one-hit mutants and double mutants are advantageous or disadvantageous (not shown).

### 3 Dynamics of mutant generation and spread

In this section we provide details of modeling generation and spread of double hit mutants, both advantageous and disadvantageous.

#### 3.1 Advantageous mutants: the time series

In the case of advantageous mutants, a reasonable measure of double mutant success is the time it takes for the double hit mutant to reach 90% of all infected cells. Figure 6 presents examples of typical infection dynamics with (right) and without (left) recombination, both for relatively low (top) and high (bottom) mutant advantage. We can see that at first, populations of single mutants rise, and at some point produce a double hit mutant, which eventually rises to domination, displacing other populations. Parameters of the system define the typical timing of this process.

In the presence of double hit mutant advantage, the following factors trade-off to determine where recombinations boost or suppress double mutant spread: (i) the constructive force of double hit mutant creation by recombinations (which enhances double hit mutant production), (ii) the destructive force of recombination that breaks down double mutants (which delays double mutant spread); and (iii) the strength of selection of the double mutant, which defines the how long the previous two factors are at play.

Consider the case where the mutant advantage ($s$) is relatively low (figure 6, top panels). For a stretch of time, two single mutant strands coexist in the population at low levels. During this period, in the absence of recombinations, a double hit mutant is created by mutations and eventually rises to domination. In the presence of recombination, double hit mutants are created at a faster rate through recombinations between single strand mutants, but since the advantage is low, they remain at low levels for a long time, which contributes to frequent breakage events through recombinations with the abundant wild type. This destructive force of recombinations is what makes recombinations delay the domination of double hit mutants.

When the mutant advantage is relatively high (figure 6, bottom panels), all the same processes take place, but their relative contributions shift. Once a double hit mutant is created, it rises relatively fast due to larger selection force, leaving the "breaking" recombinations less time to operate. On the other hand, recombinations between the two single strains still accelerate double mutant production, thus resulting in a net positive, accelerating effect of recombination on double hit mutant domination dynamics.

##### 3.2 Measuring the level of disadvantageous mutants

In the disadvantageous mutant scenario, mutant populations are less fit than wild-type viruses and do not invade. Rather, they steadily approach a balance between selection and mutation. Thus, in the long run, the double mutants converge to fluctuating around an equilibrium, the magnitude of which is determined by the mutation rate and the degree of the selective disadvantage. In the main text, a measure of double mutant relative abundance is used to assess its prominence.

To measure the relative abundance, we numerically determined the average double mutant fraction at an arbitrary time point $T = 10^5$ over many simulation runs. However, we also developed a different measure of double mutant success that yields very similar results. This measure is a temporal moving average for a single run, rather than considering averages at a fixed time point over many runs. Here, only one simulation is done at each parameter combination. The temporal moving average is calculated by (i) determining the relative fraction of cells infected with the double mutant over the total number of infected cells at each time step and then (ii) calculating the average of this fraction over the number of time steps elapsed. If the infection dies out (which only happens very rarely), results from that simulation are discarded and a new, more typical simulation is used.

The advantage of this approach is that only a single simulation is needed at each combination. The disadvantages are that (i) single simulations need to run to at least the $10^6$ time step (ii) the fraction of double mutant abundance needs to be calculated at each time step (slowing the speed of the simulation) and (iii) this measure only has relevance when a selection mutation balance is achieved. In the neutral case, when only drift occurs, this measure is not useful.

##### 3.3 Using ODEs to study selection mutation balance

As stated in the previous section, in the case of disadvantageous mutants, mutant populations are maintained by a balance between selection and mutation. It is reasonable to use ODEs to approximate this balance, and to determine whether or not recombination is a helpful force in the spread of the double hit mutant population.

The presence of recombination results in a more complex dependence between double-hit mutant creation and destruction, which can promote its spread. The key is the abundance of the single-hit mutants, compared to the abundances of double mutants and the wild types. Let us denote the abundance of single mutants of type A (or B) as at time $t$ as $Y_1$, and the abundances of double mutants and the wild types as $Y_2$ and $Y_0$ respectively. From Supplemental Figure 2, the rate of "breaking" recombinations, is

$$\rho(3/4)Y_0Y_2 + \rho(1/2)Y_1Y_2 + \rho(1/2)Y_1Y_2.$$

Similarly, from Supplemental Figure 2, the rate of "making" recombinations, is

$$\rho(1/4)Y_0Y_2 + \rho(1/4)Y_1Y_1 + \rho(1/2)Y_1Y_2 + \rho(1/2)Y_1Y_2.$$

Therefore, by setting these expressions equal to one another, we have balance at time $t$ if

$$Y_1^2 = 2Y_0Y_2. \tag{10}$$

If the right hand side of (10) is larger, then recombination between single mutants is more likely, which leads to the net creation of double mutants. If the left hand side of (10) is larger, then most recombination events will destroy the double mutant through recombination with wild-type, leading to the net loss of double mutants.

To implement these ideas, we use ODEs that do not explicitly include recombination. Instead, we look at the balance of $Y_1^2 = 2Y_0Y_2$ at time $T$ in an ODE system without recombination (thus neglecting all recombination events up to time $T$).

To define the ODE system, we assume the same 4 strains, where each strain can (forward or back) mutate at site $a/A$ and/or at site $b/B$ during each infection event. Let $x_{i,j,k,l}(t)$ be the number of cells infected with $i$ copies of wild type, $j$ copies of mutant strain A, $k$ copies of mutant strain B, and $l$ copies of the double mutant at time $t$. Let $W$, $A$, $B$, $D$ be the density of all populations infected with the wild type, mutant strain A, mutant strain B, and double hit mutant AB respectively at time $t$, that is

$$\begin{aligned} W(t) &= \sum_{0<i+j+k+l\leq N} \frac{i}{i+j+k+l} x_{i,j,k,l}(t), \\ A(t) &= \sum_{0<i+j+k+l\leq N} \frac{j}{i+j+k+l} x_{i,j,k,l}(t), \\ B(t) &= \sum_{0<i+j+k+l\leq N} \frac{k}{i+j+k+l} x_{i,j,k,l}(t), \\ D(t) &= \sum_{0<i+j+k+l\leq N} \frac{l}{i+j+k+l} x_{i,j,k,l}(t). \end{aligned}$$

Let $Z$ be the sum of all infected populations. We then have that

$$Z(t) = \sum_{0<i+j+k+l\leq N} x_{i,j,k,l}(t) = W(t) + A(t) + B(t) + D(t).$$

The wild type mutates into mutant strain A with probability $\mu(1-\mu)$, which represents mutation at point A and no mutation at point B. Similarly, the wild type mutates into mutant strain B with probably $(1-\mu)\mu$ and into the double mutant with probability $\mu^2$. This means the wild type does not mutant with probability $1-2\mu+\mu^2$. Similar mutation probabilities follow for the other strains. The ODE system is

$$x_{0,0,0,0} = \lambda(\mathcal{N}^2 - Z - x_{0,0,0,0}) - \frac{\beta}{\mathcal{N}^2} Z x_{0,0,0,0} - \frac{\gamma}{\mathcal{N}^2} Z x_{0,0,0,0} - d x_{0,0,0,0} \tag{11}$$

$$\begin{aligned} x_{i,j,k,l} &= \frac{\beta}{\mathcal{N}^2}\Big( f_W(W(1-2\mu+\mu^2) + A(\mu(1-\mu)) + B((1-\mu)\mu) + D(\mu^2))x_{i-1,j,k,l} \\ &+ f_A(W(\mu(1-\mu)) + A(1-2\mu+\mu^2) + B(\mu^2) + D((1-\mu)\mu))x_{i,j-1,k,l} \\ &+ f_B(W((1-\mu)\mu) + A(\mu^2) + B(1-2\mu+\mu^2) + D(\mu(1-\mu)))x_{i,j,k-1,l} \\ &+ f_D(W(\mu^2) + A((1-\mu)\mu) + B(\mu(1-\mu)) + D(1-2\mu+\mu^2)x_{i,j,k,l-1} - Zx_{i,j,k,l}\Big) \\ &+ \frac{\gamma}{\mathcal{N}^2}\Big( f_W(W(1-2\mu+\mu^2) + A(\mu(1-\mu)) + B((1-\mu)\mu) + D(\mu^2))x_{i-S,j,k,l} \\ &+ f_A(W(\mu(1-\mu)) + A(1-2\mu+\mu^2) + B(\mu^2) + D((1-\mu)\mu))x_{i,j-S,k,l} \\ &+ f_B(W((1-\mu)\mu) + A(\mu^2) + B(1-2\mu+\mu^2) + D(\mu(1-\mu)))x_{i,j,k-S,l} \\ &+ f_D(W(\mu^2) + A((1-\mu)\mu) + B(\mu(1-\mu)) + D(1-2\mu+\mu^2)x_{i,j,k,l-S} - Zx_{i,j,k,l}\Big) - ax_{i,j,k,l} \end{aligned} \tag{12}$$

where any population with a negative index is 0 and infection does not occur if it would result in a cell being infected with more than maximum infection multiplicity $N$ viruses.

In order to evaluate whether recombination is beneficial for a given parameter set and fitness landscape, we calculate the number of each type of virus in the system at time $T = 10^5$. If $Y_1^2 > 2Y_0Y_2$ then recombination is beneficial, otherwise it is not. The predictions from the ODE system do not perfectly match the results from the stochastic system because of many factors, including (i) the ODEs do not take into account the spatial nature of cell-to-cell transmission, (ii) the ODEs are deterministic, (iii) recombination is neglected until time $T$, and (iv) a maximum infection multiplicity $N$ must be used in order to solve the ODEs.

While they do not match perfectly, as seen in Figure 7 the ODEs do successfully qualitatively predict that for any parameter set with significant fitness difference $s$, recombination promotes double mutants under extreme negative epistasis ($\alpha$ close to 0) and suppresses them for extreme positive epistasis ($\alpha$ close to 1). This is again because for extreme positive epistasis, single mutants are less abundant, and "breaking" events dominate, resulting in recombinations suppressing double hit mutants. If on the other hand we have extreme negative epistasis, then the double hit mutants are extremely rare, and "making" events dominate, thus rendering recombination an enhancing force.

### 4 A model with a lower multiplicity of infection

The model described in the main text is characterized by a relatively high equilibrium multiplicity of infection. Figure 8 shows a typical simulation where the mean multiplicity of all infected cells is plotted as a function of time. We can see that for parameters used in figure 2 of the main text, under purely free virus transmission, a typical mean multiplicity of infection is about 4 viruses per cell, while under purely synaptic transmission, it is about 14 viruses per cell. In order to investigate how results change if infection multiplicity is lower, we designed a model with limited multiplicity. In this model, we keep track of the time, $\tau$, that has passed since the first time a cell gets infected. The longer this time, the less likely it is that the cell gets superinfected. This is because after the initial infection, the virus eventually down-regulates the receptor required for viral entry, preventing further superinfection events from occurring [1]. We assumed that after the first infection event, each subsequent infection event for a given cell can be aborted with a probability $P = 1 - e^{-\nu\tau}$, which grows to its limiting value of 1 as $\tau$ increases. In this model, under purely free virus transmission, a typical mean multiplicity of infection is just slightly over one virus per cell, while under purely synaptic transmission, it is less that 4 viruses per cell, see figure 8.

Typical results of the limited multiplicity model are presented in figure 9. The patterns that are observed for the limited multiplicity model are very similar to those reported in the main text for the basic model.

**Disadvantageous mutants.** Simulations for disadvantageous mutants are presented in panels (a,b) of figure 9, and should be compared with figure 2(c,d) of the main text.

- For disadvantageous mutants under higher infection multiplicities, recombination increased double mutant fractions at equilibrium under negative epistasis (panel (a)) and suppressed them under positive epistasis (panel (b)) if free virus transmission is dominant. This result is unchanged compared with the basic model.

- If synaptic transmission is dominant, we observed a reversal for higher multiplicities: recombination always increased double mutant fractions, even for positive epistasis. For low multiplicity, this reversal is not observed, i.e. double mutant levels are lower in the presence compared to the absence of recombination even at 100% synaptic transmission (figure 9(b)). We do find, however, that this reduction in double mutant levels in the presence of recombination is significantly less pronounced when synaptic transmission becomes more prevalent.

Hence, the conclusion that synaptic transmission protects the double mutant against the detrimental effects of recombination continues to hold under low infection multiplicities.

- The larger the percentage of synaptic transmission, the higher the level of double mutants. This is similar to the basic model, and in some sense more pronounced, as this pattern holds for near 100% synaptic transmission.

**Advantageous mutants.** Simulations for advantageous mutants are presented in panels of figure 9 (c,d,e), and should be compared with figure 2(a,b) of the main text.

- For negative and zero epistasis, recombinations play an enhancing role in double mutant spread (panels (c,d)), as in the basic model.
- For large positive epistasis and a range of fitness advantage, recombinations become detrimental for double mutant spread (panel (e)), similar to the basic model. This effect, although statistically significant, is weaker in the limited multiplicity model compared to the basic model.
- The larger the percentage of synaptic transmission, the slower the spread of mutants. This is similar to the basic model, and in some sense more pronounced, as this pattern holds over all combinations of synaptic and free virus transmission.

**Neutral mutants.** Simulations for neutral mutants are presented in panel (f) of figure 9 and should be compared with figure 3(a) of the main text.

- Recombinations result in a higher level of double mutants, as in the basic model.
- A mixture of free virus and synaptic transmission is optimal for double mutant spread, as in the basic model.

### 5 Recombinations, epistasis, and transmission mode: additional information

#### 5.1 The role of recombinations under different transmission modes

Figure 10 contains graphs supplementing Figure 4 of the main text, in the case of advantageous mutants. We show in the main text (figure 1(b) of the main text) that for advantageous mutants, under a mixture of free virus and synaptic transmission modes, recombinations mostly enhance double mutants except for a region of intermediate fitness advantages and relatively strong positive epistasis, where recombinations delay the spread of mutants.

It turns out that these results for advantageous mutants remain very similar under different mixtures of free virus and synaptic transmission. This is illustrated in figure 10, which show the time to double mutant invasion with (red) and without (black) recombinations, under free virus transmission only (a) and under synaptic transmission only (b). The time of mutant spread is presented as a function of parameter $\alpha$ (where $\alpha < 0.5$ corresponds to negative epistasis and $\alpha > 0.5$ to positive epistasis). We can see that recombination becomes disadvantageous for mutants (by delaying the time of spread) for high values of $\alpha$. In figure 1(b) of the main text, a similar outcome can be seen by looking along diagonal straight lines connecting points $\Delta_1 = A$ on the horizontal axis and $\Delta_2 = A$ on the vertical axis, where $A = |\ln(1-s)| = |\ln(1-0.05)| \approx 0.05129$.

Figure 11 contains graphs supplementing Figure 4 of the main text, in the case of disadvantageous mutants. In particular, panels (a) and (b) correspond to free virus transmission only and are similar to figure 4(a) of the main text, except they contain simulations for smaller ($s = 0.005$) and larger ($s = 0.075$) fitness disadvantage values compared to that of the main text ($s = 0.05$). Further, panels (c) and (d) correspond to synaptic transmission only ($\beta = 0$) and are similar to figure 4(b) of the main text; again, they correspond to a smaller and a larger fitness disadvantage. We observe that there is no qualitative differences for different values of fitness disadvantage.

#### 5.2 Optimal epistasis to promote double mutants

Using the results presented above and in the main text, we can investigate what level of epistasis is optimal for double mutant spread, given the transmission mode, with and without recombinations.

Figure 10 suggests that for advantageous mutants, intermediate values of epistasis ($\alpha$) are optimal to minimize the time until double mutant fixation. In order for the double mutant population to fixate past the 90% threshold, first the intermediate mutants need to overtake the wild type, and second the double hit mutant strain needs to overtake both of the intermediate mutant strains. For the mutants to quickly overtake the wild type, the intermediate mutant strains need to have a significant fitness advantage over the wild type, that is $\alpha < 1$. For the double mutant strain to quickly overtake the intermediate mutant strains, the double hit mutant needs to have a significant fitness advantage over the intermediate mutant strains, that is $\alpha > 0$. If the double hit mutant has no fitness advantage over the intermediate mutants, so $\alpha = 0$, all mutant strains have the same fitness and it takes longer for the double hit mutant strain to drift and fixate past the 90% threshold. This is true both in the absence and presence of recombination, and both under free virus and synaptic transmission. Figure 12 is a heat map showing this for a combination of free virus and synaptic transmission and a range of fitness differences $s$.

A different result is observed for disadvantageous mutants, see figure 11 and also figure 4 of the main text. As before, decreasing $\alpha \in [0, 1]$ leads to higher intermediate mutant fitness. On the other hand, decreasing $\alpha$ also leads to higher ratios of cells infected with the double hit mutant. This is because as the intermediate mutant fitness decreases, it becomes increasingly unlikely that the double hit mutant strain will be generated at all, either by mutation or recombination between the intermediate strains. Therefore, $\alpha = 0$ produces the largest level of double mutants. Again, this trend holds in the absence and presence of recombination, and both under free virus and synaptic transmission.

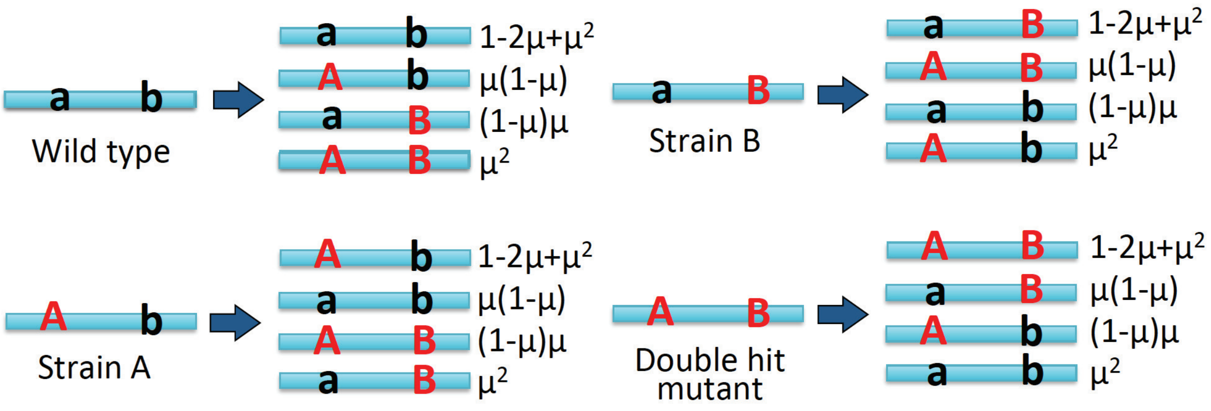

Figure 1: All mutation processes possibilities between the types, together with their probabilities.

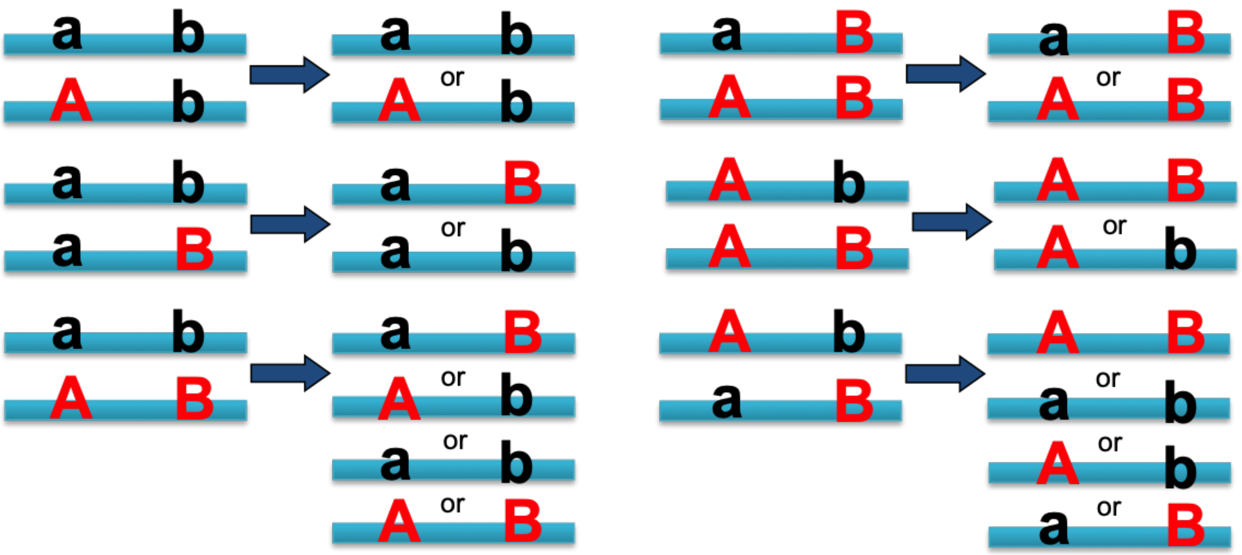

Figure 2: All recombination possibilities between the types. If the two strands are of the same strain then recombination is trivial.

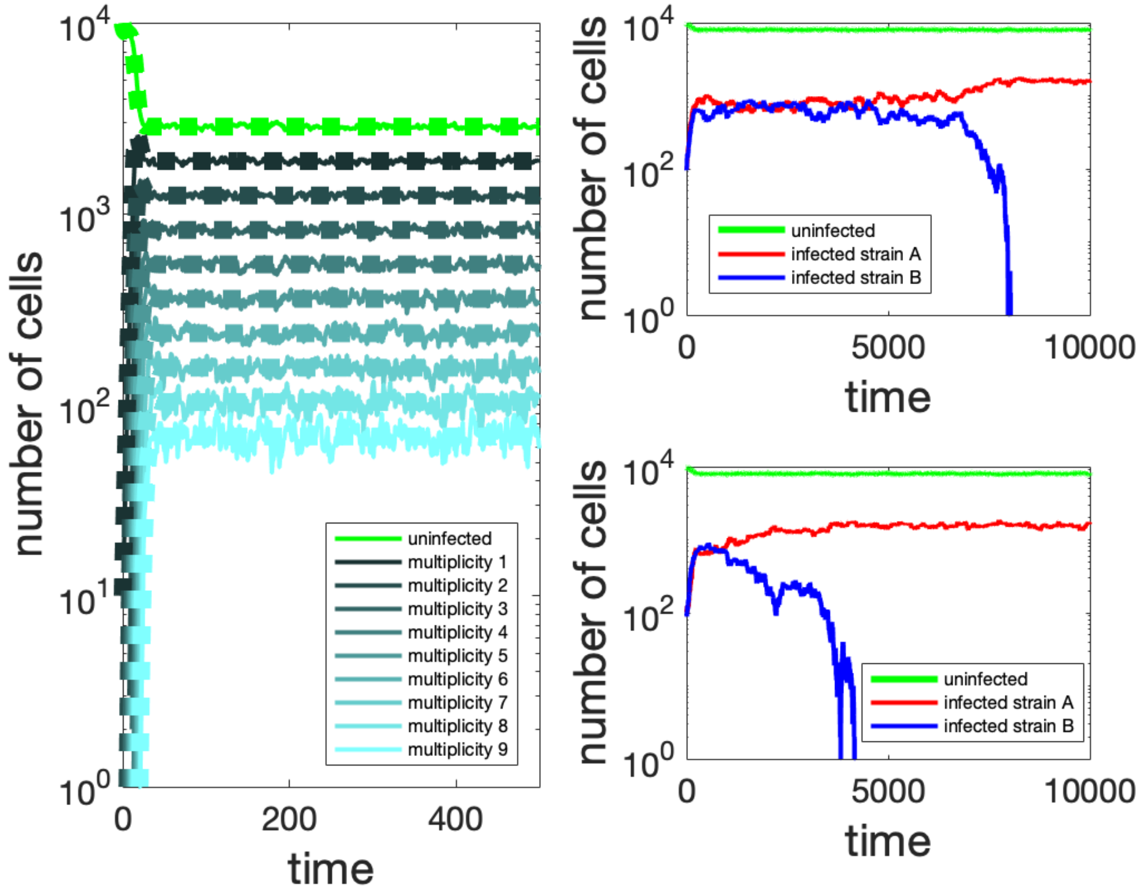

Figure 3: Left: comparison of the ordinary differential equation and agent based models for infection with a single strain. The numerical solutions for the ordinary differential equations match the agent based simulation and equilibrium values for each virus population. Right: comparison of agent based simulations for two neutral virus strains versus two non-neutral strains. Above: neutral virus strains with the same fitness. Below: strain A has higher fitness over strain B.

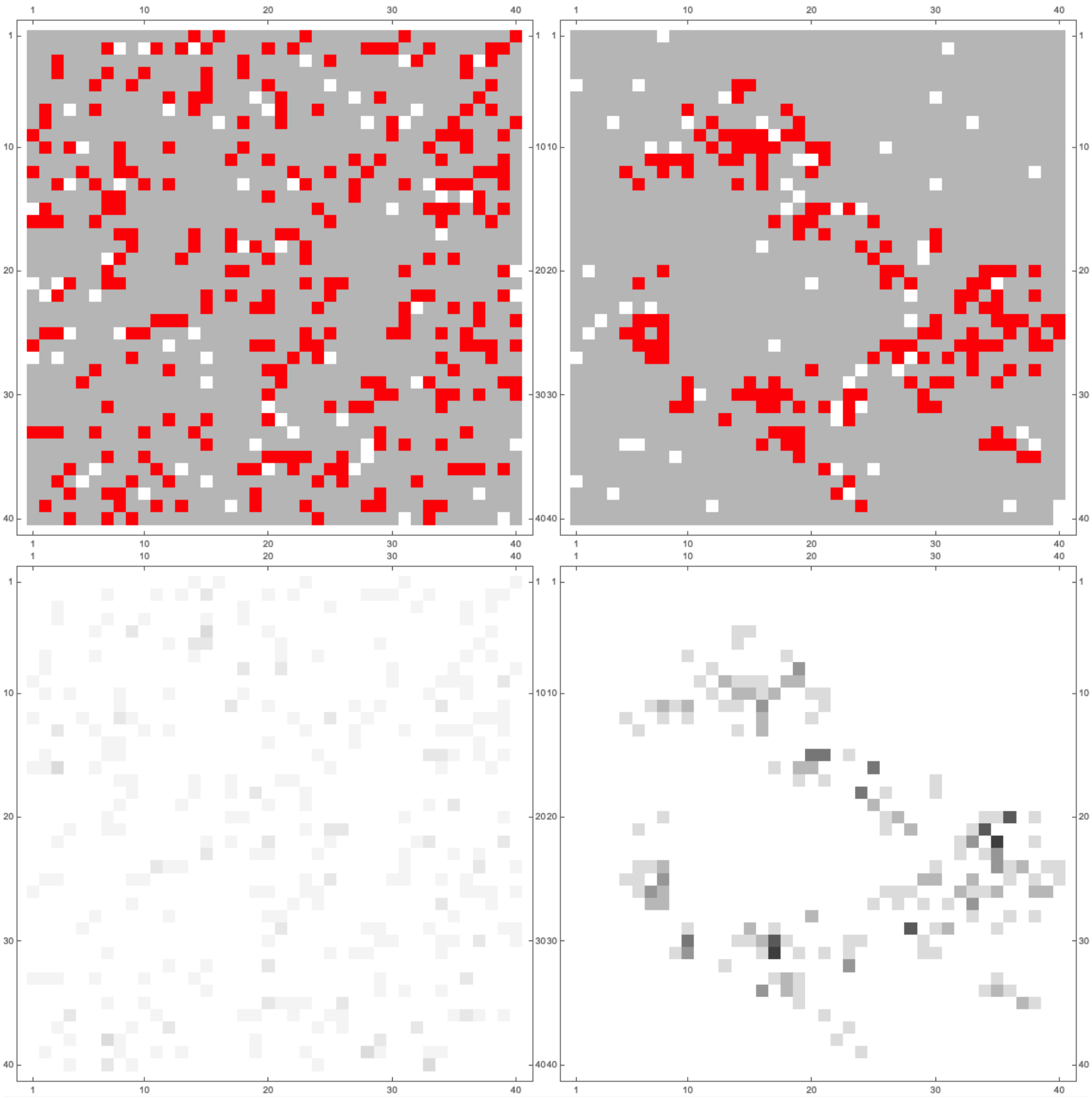

Figure 4: Example of stochastic simulations on a 40 by 40 grid at time $T = 100$. The panels on the left correspond to the same simulation with only free virus transmission. The panels on the right correspond to the same simulation with only synaptic cell-to-cell transmission. The panels on top show the infection where white grid points are empty, gray grid points are uninfected cells, and red grid points are infected cells. The panels on the bottom show the number of copies of virus each cell is infected with, where darker colors represent higher numbers.

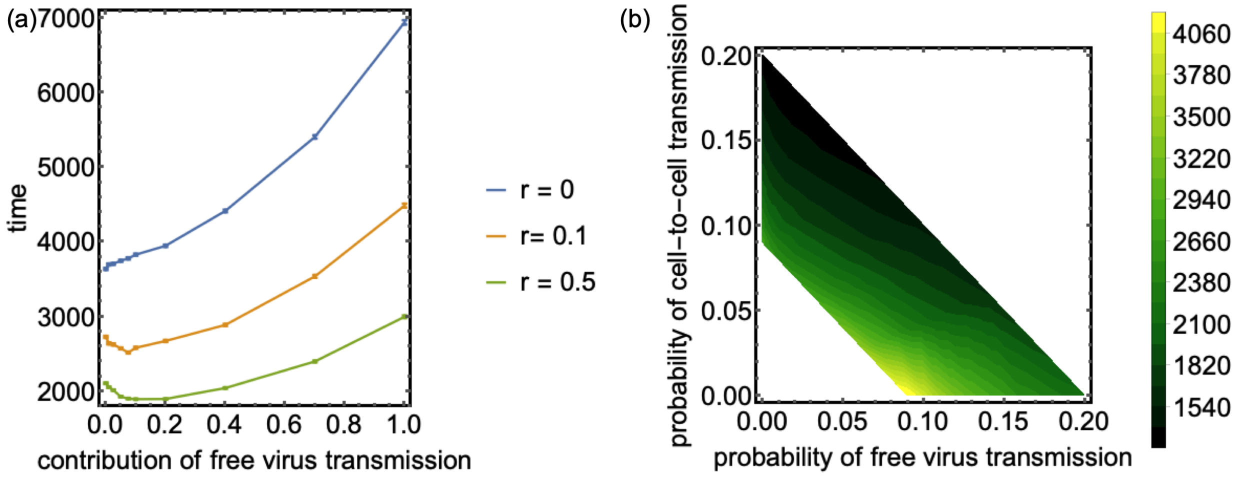

Figure 5: Generation of double mutants. (a) Time to double mutant generation, as a function of the rate of synaptic transmission, with $\beta + \gamma = 0.1$. Higher recombination rates lead to faster times to first double hit mutant generation. Standard error bars are shown. Each point represents the average over at least 12,280 runs. With a positive recombination rate $\rho \gg \mu$ a combination of free virus and cell-to-cell transmission optimizes the time to first double hit mutant generation. (b) Contour plot for the time to recombinant, plotted against the probability of free virus transmission ($\beta$) and the probability of cell-to-cell transmission ($\gamma$). Darker colors represent faster (lower) time to the creation of a recombinant virus. Diagonal lines with slope $-1$ and intercept $c$ represent fixed $\beta + \gamma = c$. For fixed $c$, a combination of both free virus ($\beta$) and cell-to-cell transmission ($\gamma$) minimizes the time to the creation of recombinant virus. Contour plot was made by running the simulations for many points on the lines with fixed $c$ for $c \in [0.09, 0.2]$. We used $\rho = .2$; enough simulations were run such that the averages with their respective standard error did not overlap. Other parameter values are $S = 3$, $\lambda = 1$, $d = 0.01$, $a = 0.02$, $\mathcal{N} = 100$, $\mu = 3 \times 10^{-5}$, and we initially infect randomly with only the wild type.

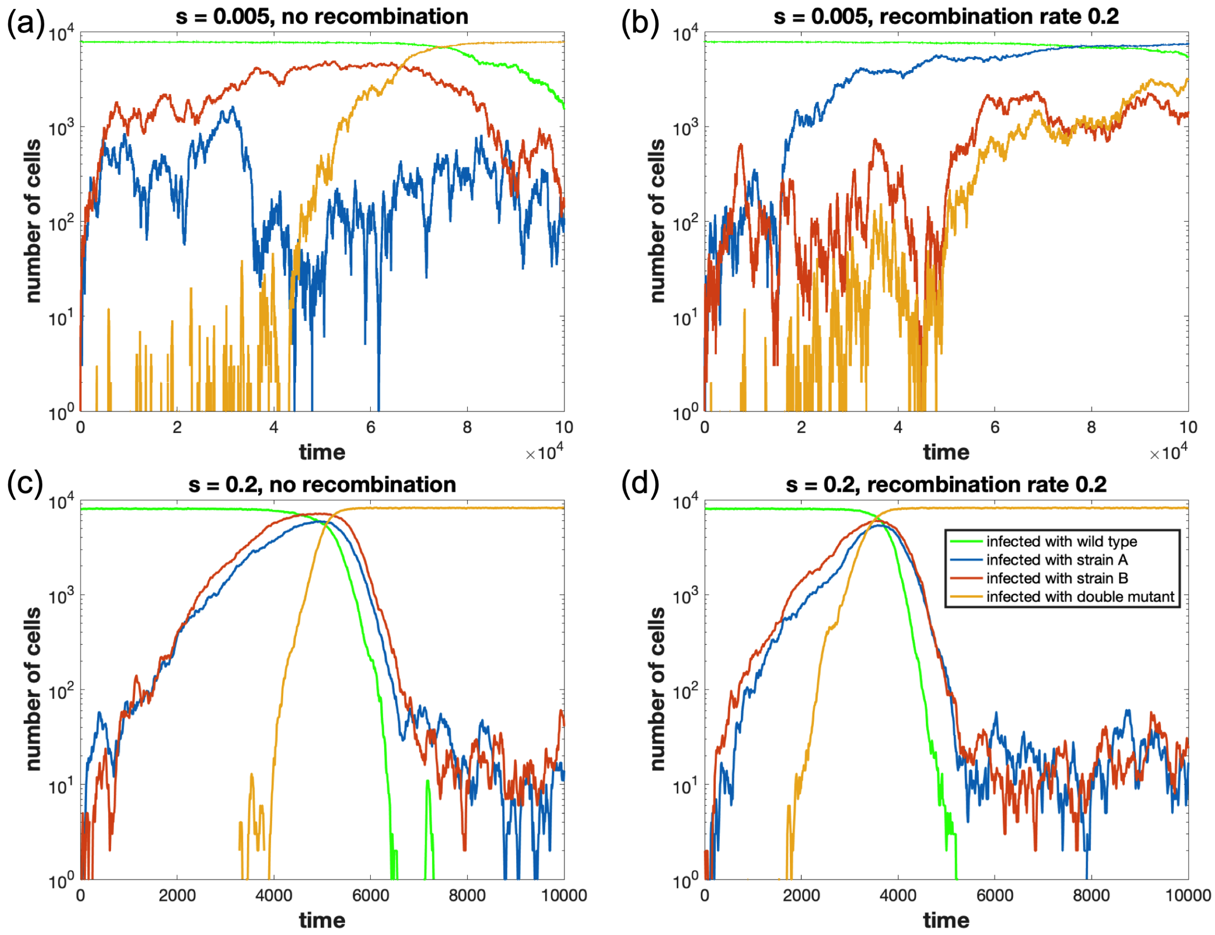

Figure 6: Advantageous mutants: the dynamics of cell populations, typical time-series. (a): slight advantage, no recombination (b): slight advantage, recombination rate $\rho = 0.2$ (c): large advantage, no recombination (d): large advantage, recombination rate $\rho = 0.2$. The parameters are: $S = 3$, $\lambda = 1$, $d = 0.01$, $a = 0.02$, $\mathcal{N} = 100$, $\mu = 3 \times 10^{-5}$, $\alpha = 0.75$, 40% free-virus transmission, and initial infection with only the wild type at equilibrium levels.

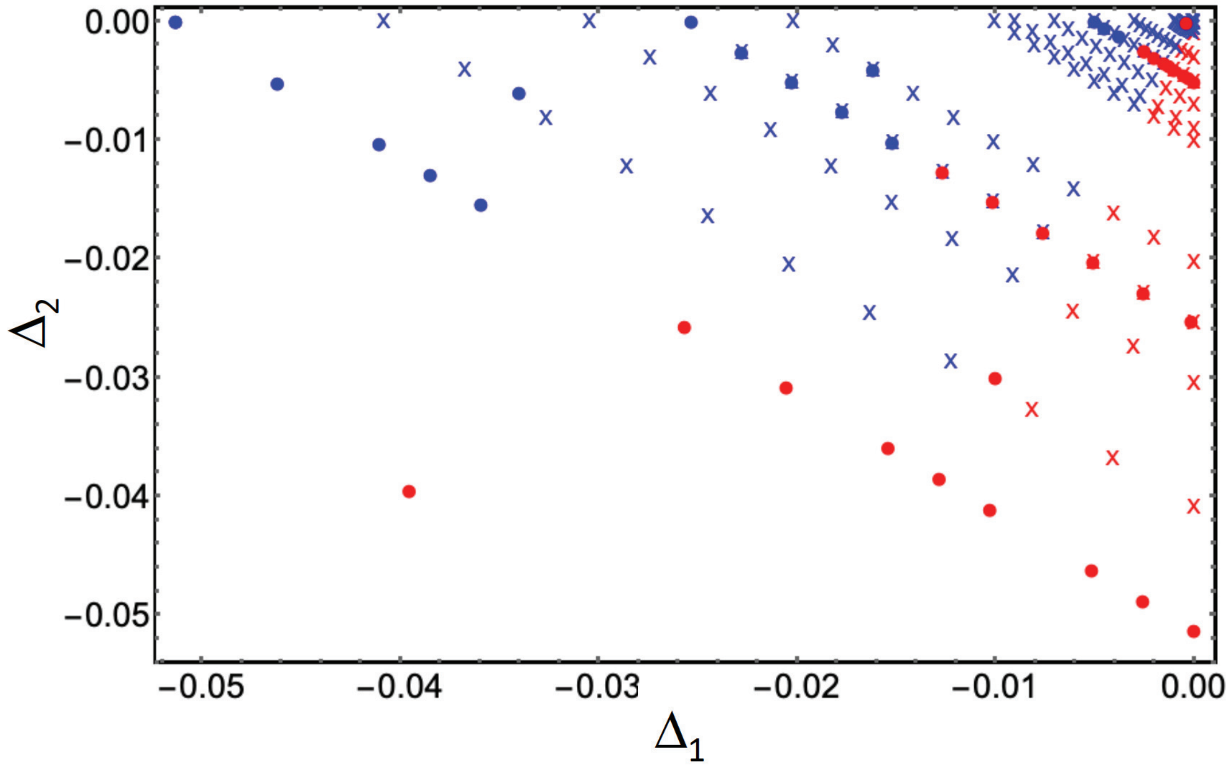

Figure 7: Results of the stochastic simulations (dots) versus the prediction from the ODE system (11-12). The horizontal axis is $\Delta_1$ and the vertical axis is $\Delta_2$, which are the relative log fitness values of single and double mutants, respectively (compare to figure 1(c) of the main text). Red (blue) corresponds to runs where double hit mutants are more (less) abundant at $T = 10^5$ in simulations with recombinations than without. We assumed 40% free-virus transmission; the rest of the parameters are $S = 3,\ \lambda = 1,\ d = 0.01,\ a = 0.02,\ \mathcal{N} = 100,\ \mu = 3 \times 10^{-5}$ and initial infection with only the wild type at equilibrium levels.

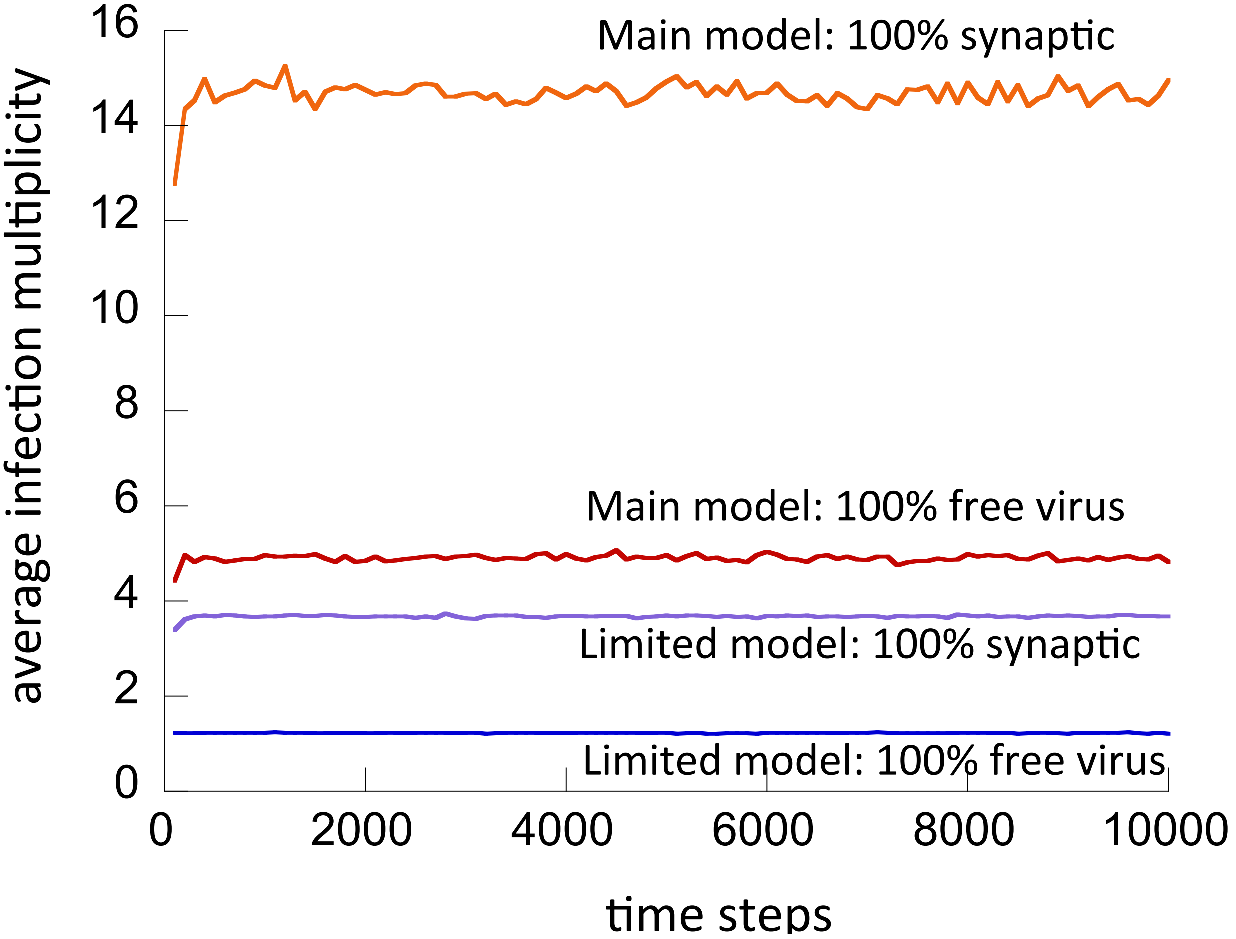

Figure 8: Mean multiplicity of infection as a function of time. The two top lines correspond to the model used in the main text (parameters as in figure 2 of the main text). The bottom two lines correspond to the limited multiplicity model, see figure 9. Results for synaptic only and free-virus only transmission are presented.

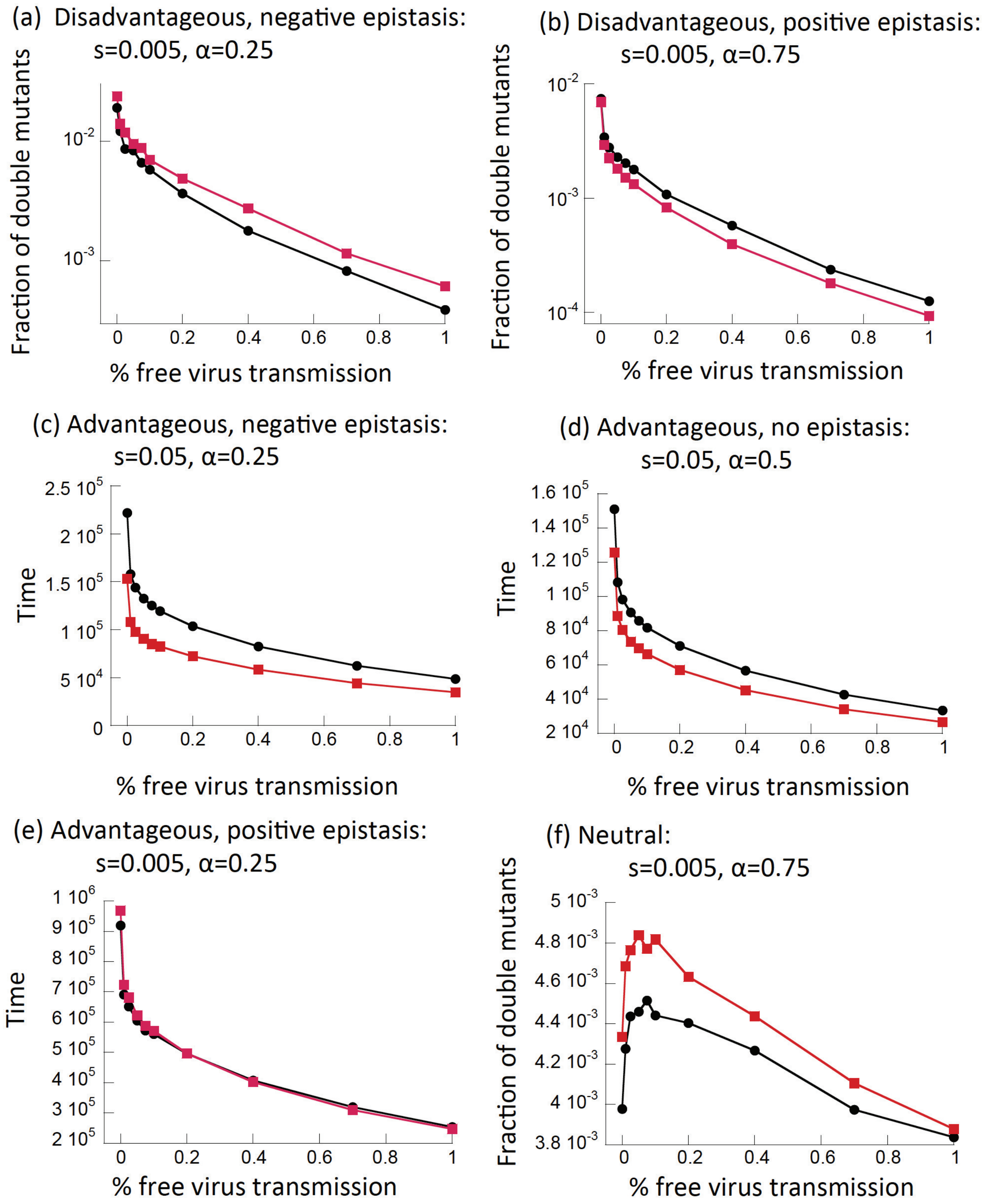

Figure 9: System with limited multiplicity: a comparison between models with (red) and without (black) recombination. (a,b) Disadvantageous mutants: the fraction of mutants (the temporal average at selection-mutation balance) as a function of the fraction of free virus transmission, under negative and positive epistasis (see figure 2(c,d) of the main text for other parameter values). (c,d,e) Advantageous mutants: the time until mutants reach 90%, for negative epistasis, no epistasis, and positive epistasis (see figure 2(a,b) of the main text for other parameter values). (f) Neutral mutants: the level of mutants as a function of the fraction of free virus transmission (see figure 3(a) of the main text for other parameter values). Vertical bars represent standard error and are too small to be seen. The additional parameter $\nu$ used in the limited multiplicity model was taken to be $\nu = 0.5$.

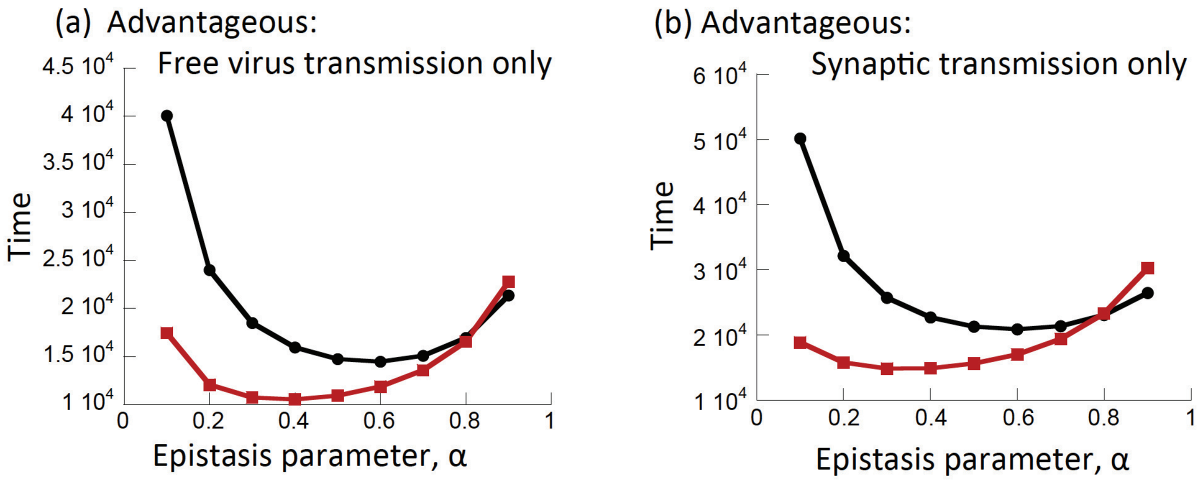

Figure 10: The role of recombinations under different transmission modes for advantageous mutants. Shown is the time for the double mutant to reach 90%. Red denotes simulations with recombination and black without recombinations. (a) Free virus transmission only, (b) synaptic transmission only. $s = 0.05$, and the other parameters are as in figure 2 of the main text. Each simulation was run at least 30,000 times. Error bars (based on standard errors) are plotted but are too small to see.

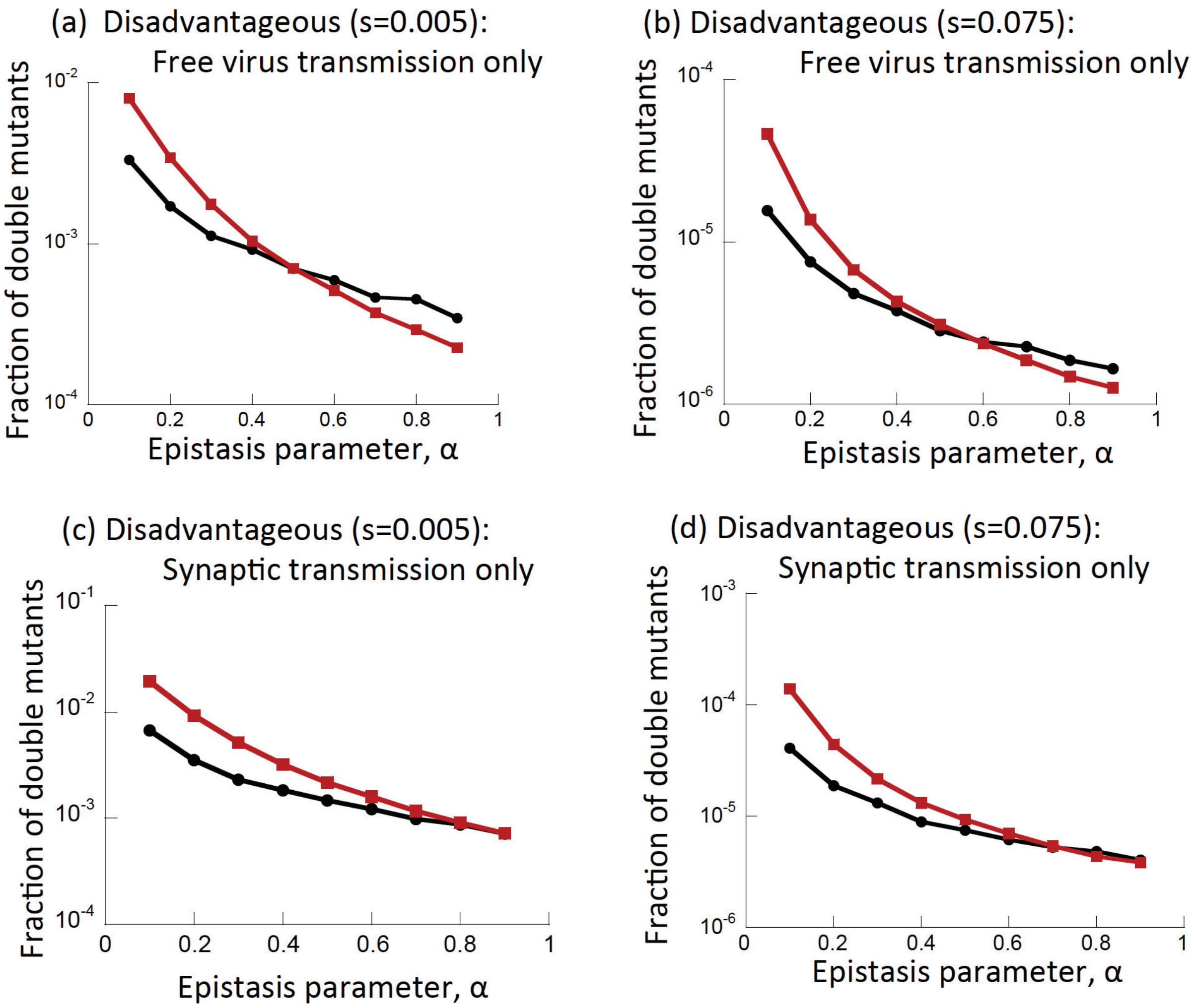

Figure 11: The role of recombinations under different transmission modes for disadvantageous mutants. Red denotes simulations with recombination and black without recombinations. (a,b) Free virus transmission ($s = 0.005$ and $s = 0.075$). (c,d) Synaptic transmission only ($s = 0.005$ and $s = 0.075$). Other parameters are as in figure 2 of the main text. The graphs show the temporal average of the double mutant at selection-mutation balance. Error bars (based on standard error) are plotted, but are too small to be visible.

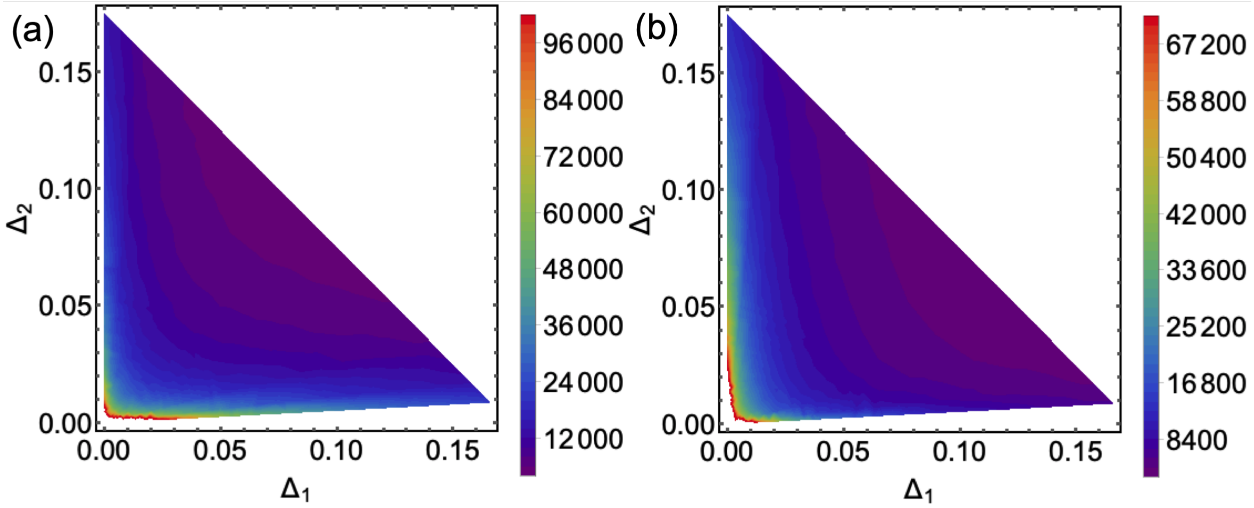

Figure 12: The optimal level of epistasis for double mutant spread. (a) Contour plot for the advantageous mutants with no recombination. The colors represent the time until double hit mutant fixation past a threshold of 90%. Lines with slope $-1$ and intercept $|\ln(1-s)|$ represent fixed $s$ with $\alpha \in [0, 1]$. Contour plots were made by running the simulations for many points on lines with fixed $s$ for $s \in [0.001, 0.16]$. The total number of points is 517. The number of simulations at each point was chosen such that the averages with and without recombination with their respective standard error did not overlap. (b) Same with recombination rate $r = 0.2$.
